# In-depth investigation of the species problem and taxonomic status of marbled crayfish, the first asexual decapod crustacean

**DOI:** 10.1101/356170

**Authors:** Günter Vogt, Nathan J. Dorn, Michael Pfeiffer, Chris Lukhaup, Bronwyn W. Williams, Ralf Schulz, Anne Schrimpf

## Abstract

The marbled crayfish is the only obligately parthenogenetic decapod crustacean and a novel research model and invasive animal on three continents. It is regarded either as a parthenogenetic form of slough crayfish *Procambarus fallax* or as a separate species named *Procambarus virginalis.* In order to investigate the species question of this unusual crayfish in detail we have identified the similarities and differences in morphology, life history, genetics, behaviour, ecology and biogeography between marbled crayfish and its most likely parent species *P. fallax.* We have investigated specimens from natural habitats, laboratory colonies and museum collections and performed a meta-analysis of our data and published data. Our *COI* based molecular tree with 27 Cambaridae confirms closest relationship of marbled crayfish with *P. fallax.* Marbled crayfish and *P. fallax* are similar with respect to morphological characters, coloration and body proportions, but differ considerably with respect to body size, fertility and longevity. The mitochondrial genes of both crayfish are similar, but ploidy level and haploid genome size are markedly different. Both crayfish are eurytopic and have two major annual recruitment periods, but marbled crayfish show different population structure and higher invasiveness. Marbled crayfish occur in tropical to cold temperate habitats of the old world, but *P. fallax* is confined to subtropical and warm-temperate habitats of the southeastern USA. Cross-breeding experiments with both crayfish revealed reproductive isolation. The application of the Evolutionary Genetic Species Concept for asexuals to all available data supports raising marbled crayfish from “forma” to species rank. A determination key is provided to discriminate *Procambarus virginalis*, the first asexual decapod species, from its parent species *P. fallax.*

## 1. INTRODUCTION

In the last 15 years, the parthenogenetic marbled crayfish has gained considerable attention in the scientific community and the public (Lukhaup, 2003; Scholtz et al., 2003; Faulkes, 2016; Scholtz, 2016; Gutekunst et al., 2018; Vogt, 2018a). It is used for research in many laboratories, is kept by aquarists worldwide and has established invasive populations in Europe, Madagascar and Japan (Chucholl, 2016). Additionally, marbled crayfish is the first crayfish with a fully sequenced genome (Gutekunst et al., 2018).

Marbled crayfish was detected in 1995 in the German pet trade and is the only obligate parthenogen among the 669 described freshwater crayfish (Crandall & De Grave, 2017) and even the 14,756 described decapod crustaceans (De Grave et al., 2009). In other higher taxa of the 66,914 Crustacea (Ahyong et al., 2011), parthenogenesis is more widespread (Martens, 1998; Dufresne, 2011; Vogt, 2018b). Earlier phylogenetic trees that included only a few crayfish species (Scholtz et al., 2003; Jones et al., 2009; Martin et al., 2010a) suggested closest taxonomic relationship of marbled crayfish to slough crayfish *Procambarus fallax* (Hagen, 1970) (Cambaridae) native to Florida and southern Georgia, which led to its provisional classification as a parthenogenetic form of slough crayfish named *Procambarus fallax* forma *virginalis* (Martin et al., 2010a).

Vogt et al. (2015) and Martin et al. (2016) presented evidence that marbled crayfish has originated from *P. fallax* by autotriploidy. Vogt et al. (2015) further established reproductive isolation in laboratory experiments and differences in life history traits and genomic features between marbled crayfish and *P. fallax*, albeit underpinned with relatively few specimens. Therefore, they suggested considering marbled crayfish a separate species and proposed the scientific name *Procambarus virginalis.* One of the authors of this paper provided later a formal description of marbled crayfish as a separate species named *Procambarus virginalis* (Lyko, 2017).

All decapod species listed in De Grave et al. (2009) are “normal” sexually reproducing species. Considering marbled crayfish as the first valid asexual species of this large crustacean group is so extraordinary and relevant that its general acceptance requires an in depth comparison of major aspects of life between marbled crayfish and its presumed parent species *P. fallax*, going far beyond the state of information presented in Vogt et al. (2015) and Lyko (2017). Of particular importance is the variability of morphological characters, life history traits and genetic markers in both crayfish that was not sufficiently considered in the previous papers. Also important is the inclusion of more *Procambarus* species in the phylogenetic tree, particularly those species that were earlier considered as closest relatives of *P. fallax* by morphological criteria (Hobbs, 1981).

In order to meet these requirements, we have compared morphology, life history, genetics, behaviour, ecology and biogeography between marbled crayfish and *P. fallax*, using numerous specimens from wild populations, laboratory colonies and museum collections. Additionally, we have performed a meta-analysis of our data and the literature data. For marbled crayfish, almost 250 publications are known, mainly based on laboratory studies (references in Faulkes, 2018; Vogt, 2018a). The number of publications on *P. fallax* is less than 50 and most of them are on its ecology (references in Hobbs, 1981; Hendrix et al., 2000; VanArman 2011; van der Heiden & Dorn, 2017; Manteuffel-Ross et al., 2018).

In the Results section we describe the similarities and differences between marbled crayfish and *P. fallax* with respect to morphological traits, coloration, body proportions, body size, fertility, longevity, mitochondrial genes, nuclear genomic features, behaviour, ecology and geographic distribution. In the Discussion section, we revisit the taxonomic position of marbled crayfish within the Cambaridae and examine the species question by applying to all data the Evolutionary Genetic Species Concept developed for asexuals by Barraclough et al., (2003) and Birky & Barraclough (2009). Asexuals are here considered as organisms that produce offspring from a single parent without fertilization, including apomictic parthenogens like marbled crayfish (Dudgeon et al., 2017).

## 2. MATERIALS AND METHODS

For this study we used data from a total of 42,536 specimens (Table 1). This number includes animals from our present experiments, unpublished earlier work of the authors, museum catalogues, GenBank and scans from published papers. Most specimens were from the wild.

**TABLE 1.** Numbers of marbled crayfish, *Procambarus fallax* and other crayfish species used for this study.

| Research field | Marbled crayfish | *P. fallax* | Other crayfish |
| --- | --- | --- | --- |
| Morphology | 64 | 50 | 0 |
| Sex determination | 1,456 | 3,902 | 0 |
| *COI*, consensus tree | 26 | 13 | 26 |
| Microsatellites | 38 | 17 | 7 |
| Body size | 1,279 | 35,388 | 0 |
| Fertility | 124 | 91 | 0 |
| Longevity | 49 | 6 | 0 |

Included are all specimens used for our meta-analysis: the experimental animals of this study, animals from unpublished datasets of the authors, and specimen from museum collections, GenBank and literature scans.

### 2.1 Animals and living conditions

Wild marbled crayfish were sampled in Lake Moosweiher (Freiburg, Germany) by hand fishing nets and baited crayfish traps. Laboratory raised marbled crayfish came from the colonies of G.V. and N.J.D. and museum specimens came from the collection of the North Carolina Museum of Natural Sciences (NCSM, Raleigh, USA). Wild slough crayfish *Procambarus fallax* (Hagen, 1870) were caught in the Everglades (Florida, USA) with throw traps and laboratory raised *P. fallax* were obtained from the colonies of N.J.D. and G.V. Additional specimens came from the collections of the NCSM and the Smithsonian Institution National Museum of Natural History (USNM, Washington, USA). Our crayfish colonies were reared in compliance with the Directive 2010/63/EU of the European Union, the US National Research Council’s Guide for the Care and Use of Laboratory Animals and institutional standards.

Lake Moosweiher, the sampling site of marbled crayfish, is a mesotrophic gravel pit located in the outskirts of the city of Freiburg, Germany (N48.031511, E7.804962E), at 218 m above sea level. It has an area of 7.6 ha and a maximum depth of 8 m. The lake is fed by ground water and drained by a brook that is also inhabited by marbled crayfish. Water temperatures reach a maximum of 26°C in summer and drop down to about 5°C in winter. In strong winters, the lake is ice-covered for shorter periods of time (Chucholl & Pfeiffer, 2010). The water is slightly alkaline (pH 7.1-8.1), and conductivity is relatively low ranging between 299 and 352 μS cm^−1^. Visibility depth is between 2.5 m and 5.5 m. Oxygen concentration is 7.6-11.7 mg L^−1^ at the surface and 2.8-5.3 mg L^−1^ at the bottom, depending on season. The riparian habitat is tree lined (mainly black alder *Alnus glutinosa*) and the littoral zone is dominated by tree roots, larger stones, patches of common reet *Phragmites australis*, and gravelly, sandy and muddy areas. The lake provides different types of food resources suitable for crayfish including bivalves, gastropods, insects, fishes, aquatic macrophytes (e.g. *Potamogeton sp.* and *Elodea canadensis*) and terrestrial plant detritus. Crayfish predators like catfish (*Silurus glanis*), pike (*Esox lucius*), perch (*Perca fluviatilis)* and eel (*Anguilla anguilla*) are abundant. There are also some waterfowl like Great Crested Grebe (*Podiceps cristatus*) that prey on crayfish. Moreover, marbled crayfish shares the lake with spiny-cheek crayfish *Faxonius limosus* (Rafinesque, 1817) (former name: *Orconectes limosus*) (Chucholl & Pfeiffer, 2010; Günter, 2014).

The wild *P. fallax* used for this study came from the Everglades in southern Florida, a large (9,150 km) low-nutrient subtropical wetland with soft-bottom substrates and a pH close to neutral. The Everglades has a distinct wet and dry season and water depths typically range spatially and temporally from 0.1-1.2 m. *P. fallax* migrate among wetland habitats in response to the rising and falling water (Cook et al., 2014; van der Heiden & Dorn, 2017). In the Everglades, *P. fallax* is omnivorous and readily feeds on animals like gastropods (Dorn, 2013), but bears a trophic signature closer to an herbivore-detrivore (Sargeant et al., 2010). The densities are commonly 1-10 specimens per m^2^ and are limited, in large part, by predatory fishes (Dorn & Cook, 2015). *P. fallax* is also prey for wildlife like alligators and otters and in particular, they are dominant contributors to the nestling diet of the most abundant wading bird, the White Ibis *Eudocimus albus* (Boyle et al., 2014).

The marbled crayfish and *P. fallax* of the indoor colonies of G.V. were raised individually or communally in simple settings and fed with TetraWafer mix pellets throughout life as described earlier (Vogt, 2008a). Tap water (pH 7.5) was used for rearing. Water temperature fluctuated between 25°C in summer and 15°C in winter and photoperiod was natural. The founder specimen of the marbled crayfish colony started in 2003 was obtained from biologist Frank Steuerwald who has kept this crayfish in purebred line since 1995, the year of detection of the marbled crayfish. The founder pair of the *P. fallax* colony was obtained from the German pet trade in February, 2014. The founder specimens of the colony of *P. fallax* of N.J.D originated from the Everglades, southern Florida (N26.070194, W80.677097) and were maintained in self-sustaining populations outdoors in multiple large 500 L (1.1 m^2^) round mesocosms with shelters and natural vegetation/algae and invertebrates (e.g., snails). They fed weekly with shrimp pellets (35% protein), but also fed on the available algal and invertebrate resources.

### 2.2 Morphological characters, coloration and body proportions

Morphological characters were compared by the same person (G.V.) between 30 marbled crayfish of 38.1-101.1 mm TL from Lake Moosweiher and the laboratory colony of G.V. and 16 *P. fallax* of 45.0-69.1 mm TL from the Everglades and the colonies of N.J.D and G.V. Photographic documentation was done with a Canon EOS 7D und a Sigma 105 close-up lens. Coloration of both crayfish and its variability was documented for numerous wild and laboratory reared specimens as well. Body proportions were determined for 37 marbled crayfish from Lake Moosweiher, 22 marbled crayfish from the colony of G.V., 5 marbled crayfish from the collection of the NCSM, 39 *P. fallax* from the NCSM and USNM collections coming from various habitats in Florida and Georgia, 7 *P. fallax* from the Everglades and surrounding wetlands and 4 *P. fallax* from the colony of N.J.D. Measured were the ratios of total length to carapace length (TL: tip of rostrum to end of telson; CL: tip of rostrum to end of carapace), CL to postorbital carapace length (POCL: orbit to end of carapace), CL to carapace width (CW: widest part of carapace) and CL to areola length (AL). These data were measured with a calliper to the nearest 0.1 mm.

### 2.3 Body size, fertility and longevity

The body size data for marbled crayfish were inferred from 172 specimens of the colony of G.V., 37 specimens from Lake Moosweiher and further 1070 specimens from literature sources: 496 from Lake Moosweiher (Chucholl & Pfeiffer, 2010; Günter, 2014; Lehninger, 2014; Wolf, 2014; Buri, 2015), 66 from wetlands of Madagascar (Jones et al., 2009: Kawai et al., 2009), 39 from thermal Lake Hévíz in Hungary (Lőkkös et al., 2016), 450 from Lake Šoderica in Croatia (Žižak, 2015; Cvitanić, 2017) and 27 from Chorvátske rameno canal in Slovakia (Lipták et al., 2017). Corresponding data on *P. fallax* came from unpublished field data of 35,388 specimens sampled in the Everglades by N.J.D., Joel Trexler (Florida International University, Miami, FL) and Dale Gawlik (Florida Atlantic University, Boca Raton, FL) and 39 specimens of the NCSM and USNM collections.

Fertility (eggs per female and clutch) of marbled crayfish was calculated from 124 females coming from the laboratory populations of G.V. and N.J.D. and published datasets from the laboratory (Seitz et al., 2005) and wild populations in Madagascar (Jones et al., 2009) and Slovakia (Lipták et al., 2017) (Table 2). The published egg count and carapace length data were digitized from the respective figures using Image J software (v. 1.51j8). These data were statistically compared to fertility of 91 *P. fallax* females, mostly coming from collections in the Everglades by N.J.D. and J. Trexler and published data from Hendrix et al. (2000) (Table 2). Fertility comparisons were conducted using analysis of covariance (type III SS) with female carapace length (mm) as the covariate using the R (v. 3.4.0) software package.

**TABLE 2.** Datasets accessed for comparison of fertility between marbled crayfish and *Procambarus fallax.*

| Animals | n | Environment | Location | Source |
| --- | --- | --- | --- | --- |
| Marbled crayfish | 58 | Wetlands, ditches | Madagascar | Jones et al., 2009 |
| Marbled crayfish | 11 | River | Slovakia | Lipták et al., 2017 |
| Marbled crayfish | 32 | Laboratory | Germany | Seitz et al., 2005 |
| Marbled crayfish | 16 | Laboratory | Germany | Vogt, unpublished |
| Marbled crayfish | 4 | Lake | Germany | Vogt, unpublished |
| Marbled crayfish | 3 | Laboratory | Florida | Dorn, unpublished |
| *P. fallax* | 59 | Wetlands | Florida | Trexler, unpublished |
| *P. fallax* | 14 | Wetlands | Florida | Hendrix et al., 2000 |
| *P. fallax* | 13 | Experimental wetlands | Florida | Dorn, unpublished |
| *P. fallax* | 5 | Various | Florida | Williams, unpublished |

Listed are habitats and sample sizes (n) of crayfish for which we could obtain both carapace lengths and egg counts.

Estimates on the longevity of marbled crayfish and *P. fallax* are based on unpublished and published (Vogt, 2010) records from the laboratory populations of G.V. and size-frequency analyses of wild populations in Lake Moosweiher (Wolf, 2014) and the Everglades (van der Heiden & Dorn, 2017).

### 2.4 Genetic analyses

For the investigation of the taxonomic relationship of marbled crayfish with *P. fallax* and other Cambaridae, we sequenced fragments of 606 bp of the mitochondrial *cytochrome oxidase subunit 1* (*COI*) genes of 4 marbled crayfish from Lake Moosweiher, 3 marbled crayfish from the NCSM and 4 *P. fallax* from the laboratory population of N.J.D. DNA was purified from muscle tissue of the legs using the “High Salt DNA Extraction Protocol for removable samples” (Aljanabi & Martinez, 1997). Polymerase chain reaction (PCR) was used to amplify the *COI* fragment using primers LCO1490 and HCO2198 (Folmer et al., 1994). PCR was performed in 20 μl volumes containing 1x Taq buffer, 2.5 mM MgCl_2_, 0.25 mM dNTP, 0.5 μM of each primer, U Taq polymerase, ca. 50-100 ng DNA and ddH_2_O. After an initial denaturation step of 4 min at 94°C, cycling conditions were 35 cycles at 94°C for 45 s, 47°C for 45 s and 72°C for 1 min with a final elongation step of 10 min at 72°C. Sequencing of both DNA strands was done by SeqIT (Kaiserslautern, Germany).

Our *COI* sequences and sequences of 26 further freshwater crayfish species downloaded from GenBank (Table 3) were then used for phylogenetic tree construction. All sequences were aligned with Geneious. jModelTest (Darriba et al., 2012) was used to estimate the best nucleotide substitution model and the HKY+G model was selected by Bayesian information criterion (BIC). Phylogenetic relationships were analysed using MrBayes 3.2.6 (Ronquist & Huelsenbeck, 2003). Four independent chains of 10 million generations were run with a subsample frequency of one thousand after a burn-in period of 1 million.

**TABLE 3.**
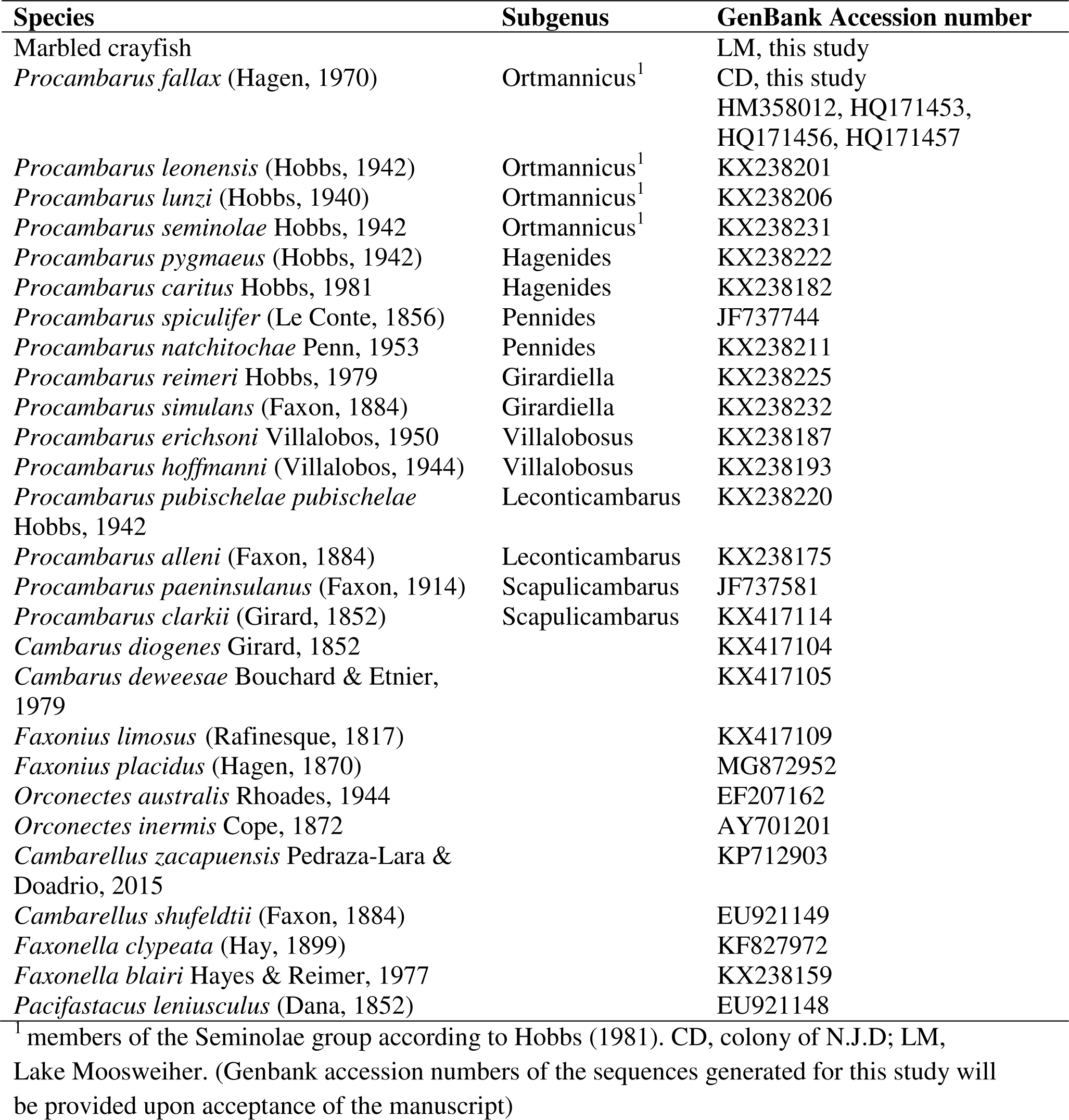
Mitochondrial COI sequences used for phylogenetic analysis.

**TABLE 4.** Body proportions of marbled crayfish and *Procambarus fallax.*

| **Body proportion** | **Marb. crayfish colony G.V.**<br>n=22<br>30.1-86.1 mm | **Marb. crayfish Moosweiher**<br>n=37<br>26.7-101.1 mm | ***P. fallax* Everglades**<br>n=11<br>45.0-69.1 mm | ***P. fallax* museums**<br>n=39<br>31.9-88.9 mm |
| --- | --- | --- | --- | --- |
| TL:CL | 2.26±0.048<br>r=1.95-2.36 | 2.16±0.047<br>r=2.08-2.25 | 2.16±0.069<br>r=2.05-2.30 | 2.12±0.048<br>r=2.00-2.19 |
| CL:CW | 2.01±0.049<br>r=1.94-2.09 | 2.31±0.137<br>r=2.10-2.61 | 2.39±0.319<br>r=1.89-2.81 | 2.33±0.232<br>r=2.04-3.09 |
| CL:POCL | 1.25±0.030<br>r=1.19-1.31 | 1.39±0.060<br>r=1.28-1.54 | 1.34±0.059<br>r=1.21-1.42 | n.d. |
| CL:AL | 2.70±0.089<br>r=2.56-2.83 | 3.07±0.215<br>r=2.64-3.47 | 3.20±0.225<br>r=2.76-3.56 | n.d. |

Figures in header row give the numbers of specimens measured (n) and their TL range. Figures in table are means ± standard deviation and ranges (r). AL, areola length; CL, carapace length; CW, carapace width; POCL, postorbital carapace length; TL, total length; n.d., not determined.

The nuclear microsatellites *PclG-02, PclG-04* and *PclG-26* (Belfiore & May, 2000) were analysed for the same marbled crayfish and *P. fallax* as used for *COI* analysis to test their diagnostic value for species discrimination as suggested earlier (Vogt et al., 2015). The conditions for PCR were as follows: DNA was denatured at 95°C for 2 min, followed by 35 cycles of denaturation at 95°C for 30 s, and annealing at 65°C for 30 s for *PclG-02* and 55°C for 30 s for *PclG-04* and *PclG-26*. Then followed an elongation step at 72°C for 1 min and final elongation at 72°C for 5 min. Fragment analysis was performed on a Beckman Coulter CEQ 8000 eight capillary sequencer using the Beckman Coulter DNA Size Standard Kit 400 bp. Loci were scored with GeneMarker, v.2.6 (SoftGenetics, State College, PA, USA).

### 2.5 Behaviour, ecology and biogeography

Behavioural, ecological and biogeographic data on marbled crayfish and *P. fallax* came from our unpublished and published records and the literature and were compiled to current overviews. The behavioural data were mainly collected from the laboratory colonies of G.V. The ecological data were collected in Lake Moosweiher for marbled crayfish and in the central Everglades for *P. fallax.* Earlier field work of authors of this paper has contributed to the distribution maps of marbled crayfish and *P. fallax.* M.P. detected the marbled crayfish population in Lake Moosweiher in 2009, which was the first self-sustaining population in Europe (Chucholl & Pfeiffer, 2010), N.J.D. worked on various wetlands of southern Florida (e.g., Dorn & Volin, 2009; van der Heiden & Dorn, 2017) and found *P. fallax* further south than described by Hobbs (1981) and Crandall (2010), and C.L. performed several crayfish surveys in southeastern USA including all of Florida and Georgia (Lukhaup, 2003; unpublished data). The references used for the meta-analyses on behaviour, ecology and biogeography are listed in the respective Results sections.

### 2.6 Author contributions

G.V. conceived and coordinated the study, made the morphological, ecological and biogeographic comparisons and contributed to the life history analysis. N.J.D. carried out the analysis of the life history data, particularly the fertility statistics, participated in the ecological comparison, sampled *P. fallax* tissues for genetic analysis and provided specimens of *P. fallax* for morphological comparison and information on the biology of *P. fallax.* M.P. sampled the marbled crayfish from Lake Moosweiher, provided data on its ecology in this lake, and sampled tissues for genetic analysis. C.L. made the photographic documentation of the taxonomically relevant morphological traits and provided information on *P. fallax* in Florida and Georgia. B.W.W. measured the body proportions of the museum samples of *P. fallax* and provided tissue samples for genetic analysis. A.S. and R.S. performed the genetic analyses and calculated the phylogenetic tree. G.V. wrote the draft manuscript and all authors contributed to the manuscript and edited the final version.

## 3. RESULTS

### 3.1 Morphology of marbled crayfish and comparison with *Procambarus fallax*

In this section we present data on morphological characters, coloration and body proportions of marbled crayfish and *P. fallax.* The morphology of marbled crayfish is described and illustrated in detail because it is not yet sufficiently covered in the literature. Hobbs (1981) described *P. fallax* in detail and considered it an extremely variable species. The same holds for marbled crayfish despite its genetic uniformity. Therefore, special emphasis is given to trait variability in both crayfish.

#### 3.1.1 Habitus and taxonomically relevant morphological characters

Comparison of 30 females of marbled crayfish of 38.1-101.1 mm TL and 16 females of *P. fallax* of 45.0-69.1 mm TL revealed that the habitus of the females of both crayfish is similar (Fig. 1a, b, d). Both crayfish are relatively slender cambarids. The maximum body size (TL) is ~11 cm in marbled crayfish and ~9 cm in *P. fallax* (see 3.2.2), corresponding to maximum fresh weights of approximately 35 g and 22 g, respectively. Marbled crayfish have only females, whereas *P. fallax* have females and males. The males of *P. fallax* (Fig. 1c) have different body proportions than the females and possess larger and more robust chelipeds (Fig. 1b). The taxonomically relevant morphological characters (defined in Hobbs, 1972, 1981) are similar in marbled crayfish and *P. fallax* females as well and vary considerably in both crayfish. The following description and illustration of these characters is mainly for marbled crayfish but also applies for *P. fallax.* It is guided by earlier diagnoses for marbled crayfish (Kawai et al., 2009; Lyko, 2017) and *P. fallax* (Hobbs, 1981).

**FIGURE 1:**
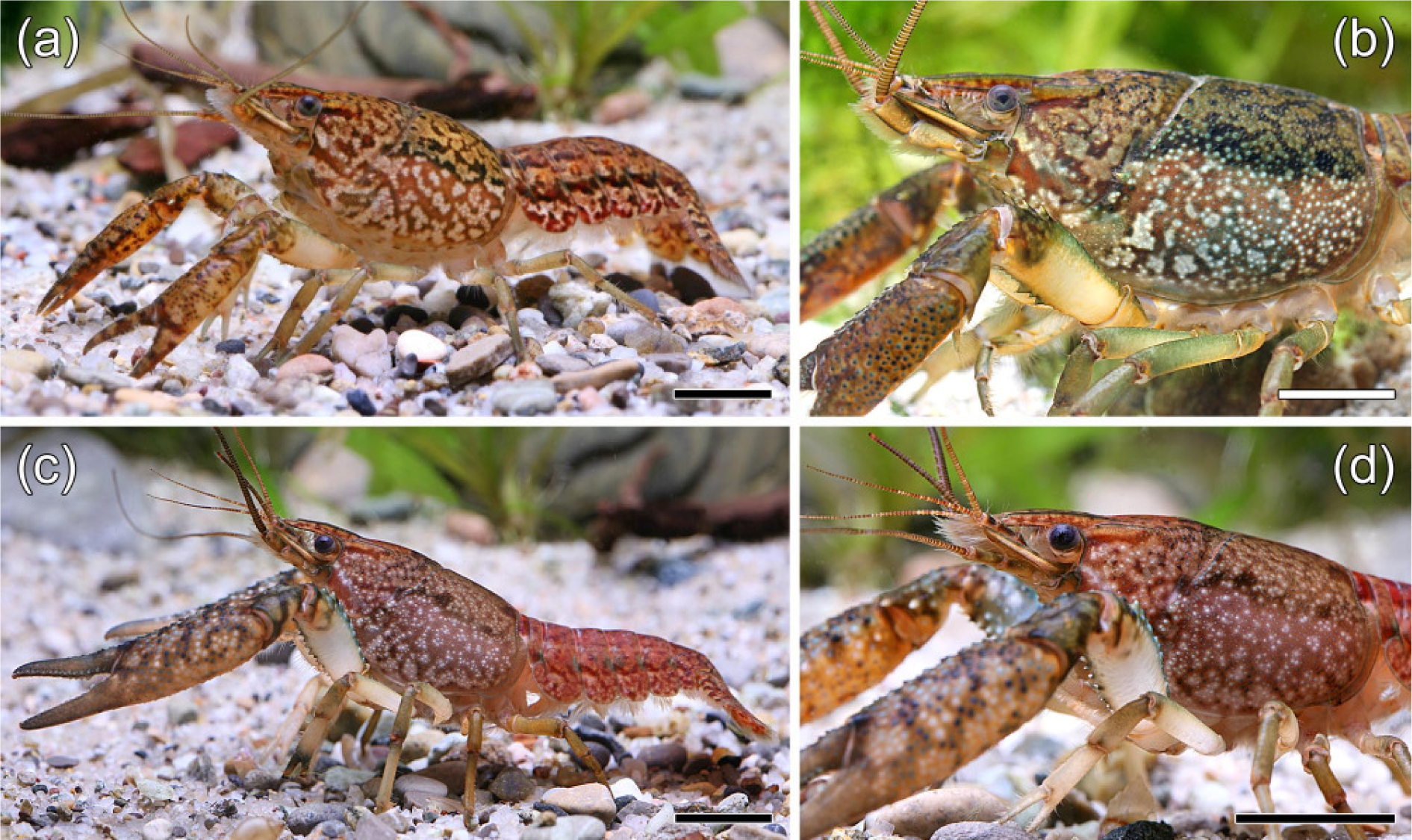
Comparison of the habitus of marbled crayfish and *Procambarus fallax.* (a) Laboratory-raised marbled crayfish female. Bar: 1 cm. (b) Cephalothorax of wild marbled crayfish. Bar: 1 cm. (c) Laboratory-raised male of *P. fallax.* Note large chelipeds. Bar: 1 cm. (d) Cephalothorax of laboratory-raised *P. fallax* female. Bar: 1 cm.

The eyes of marbled crayfish are well-developed and pigmented (Figs. 2a, 3a). The body is also well pigmented showing variable patterns of lighter spots on darker background (Figs. 1a, 2b, 6a, b, d, e). The antenna is longer than the body if not regenerated, and the pleon inclusive of the telson is longer than the carapace (Fig. 2a).

**FIGURE 2.**
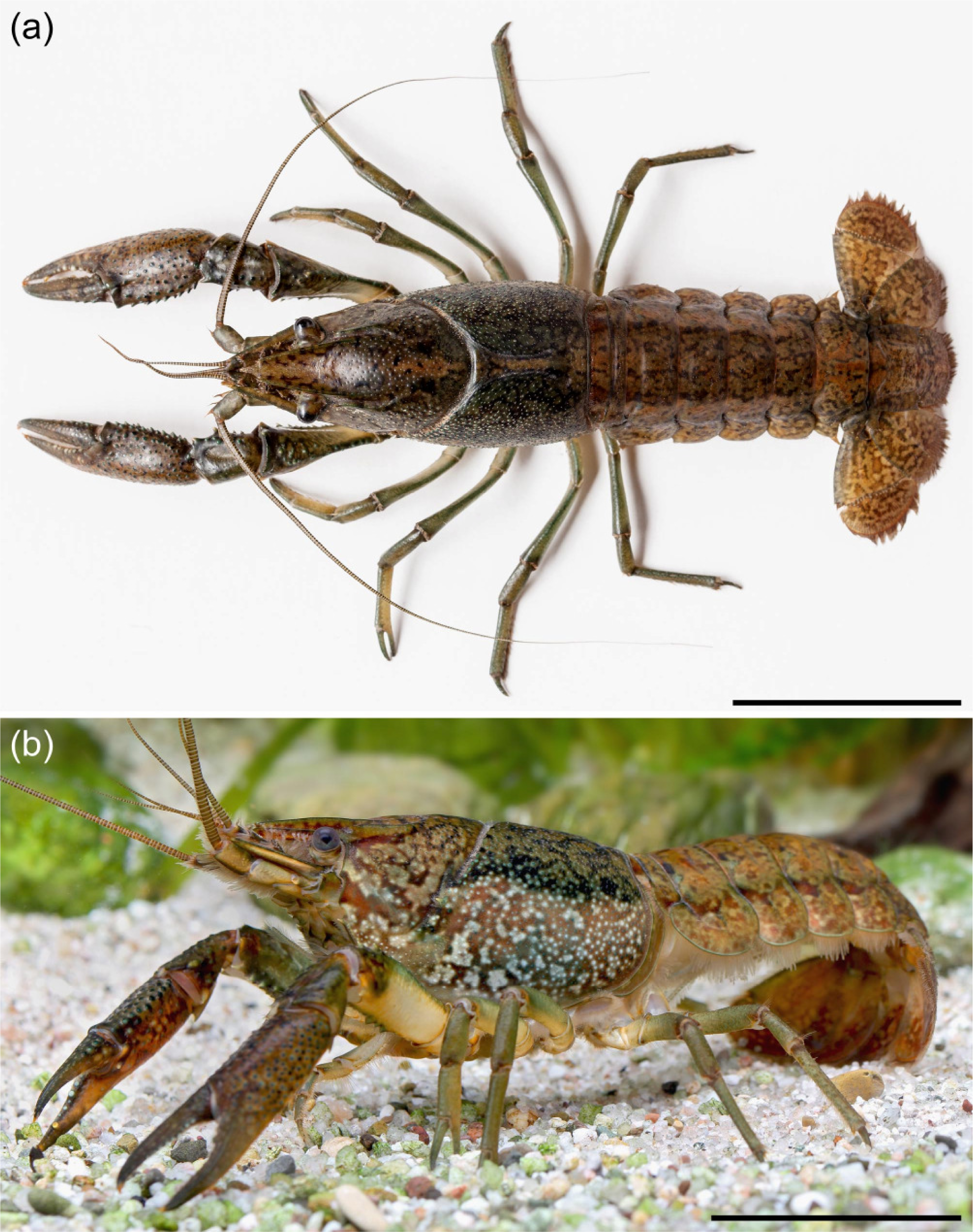
Marbled crayfish from Lake Moosweiher. (a) Dorsal view of specimen of 9.3 cm TL. Bar: 3 cm. (b) Lateral view of specimen of 10.1 cm TL. Bar: 3 cm.

**FIGURE 3.**
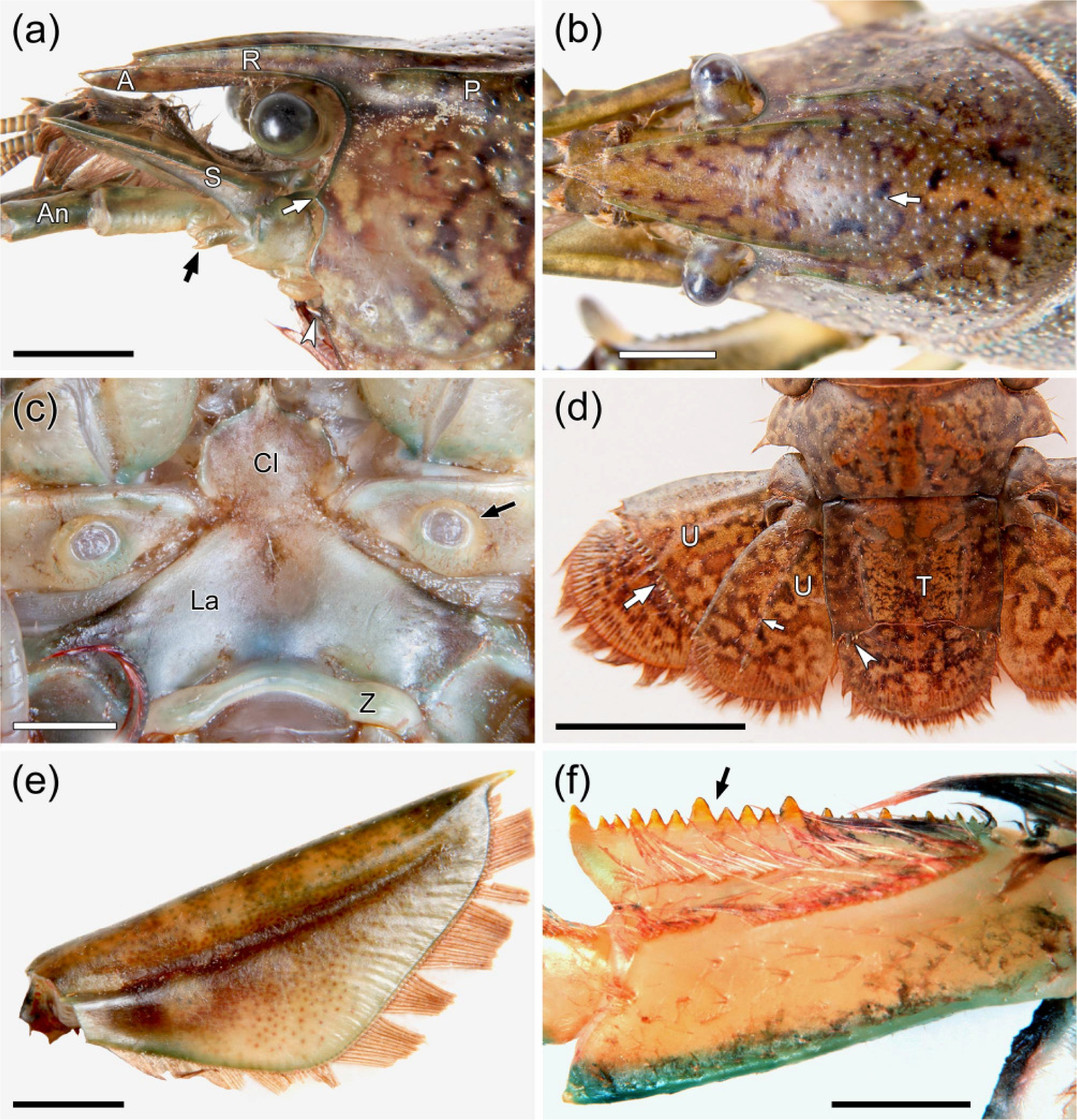
Taxonomically relevant characters on body of marbled crayfish of 9.3 cm TL. (a) Side view of cephalic region with eye, antennae (An), rostrum (R) with acumen (A), postorbital ridge (P) and scaphocerite (S). Black arrow, spine on ischium of antenna; white arrow, suborbital angle; arrowhead, branchiostegal spine. Bar: 5 mm. (b) Dorsal view on rostrum. Arrow, surface depressions with sensory setae. Bar: 5 mm. (c) Epistome composed of cephalic lobe (Cl), lamella (La) and zygoma (Z). Arrow, opening of antennal gland. Bar: 2 mm. (d) Dorsal view of telson. Small and large arrows denote rows of spines and tubercles on uropod rami, and arrowhead denotes mediolateral group of spines on telson. Bar: 10 mm. (e) Dorsal view of scaphocerite. Bar: 2 mm. (f) Inner side of ischium of 3rd maxilliped with row of teeth (arrow) on mesial margin. Bar: 1 mm.

The carapace of marbled crayfish is dorsally slightly bulged (Fig. 1b, 2b) and is broadest at two thirds of its length (Fig. 1a). It is slightly higher than wide. The suborbital angle is not acute (Fig. 3a). The dorsal surface is equipped with small punctuate depressions with sensory setae (Fig. 3b) and the lateral surface of the carapace shows granular protuberances (Fig. 6b).The carapace is studded with one acute cervical spine and one acute branchiostegal spine on either side (Fig. 3a). The rostrum has a central spiky acumen and elevated lateral margins with spines at their anterior ends (Fig. 3a, b). A median carina is lacking. The postorbital ridge is well developed and has a cephalic spine (Fig. 3a). The areola (Fig. 6a) constitutes approximately one third of the carapace length and is on average 5-6 times as long as wide.

The first antenna lacks a conspicuous fringe on the mesial border and has a spine on the ischium (Fig. 3a). The antennal scale (scaphocerite) is approximately 2.5 times as long as wide and is widest at mid-length. Its lateral margin is thickened and terminates in a spine (Fig. 3e). The epistome has a circular cephalic lobe that bears a cephalo-median projection (Fig. 3c). The lateral margin of the lobe is thickened and has no setae. The lamellae are plane and have truncate latero-posterior corners. The zygoma is slightly arched (Fig. 3c). The ischium of the 3rd maxilliped bears a row of teeth on the mesial margin (Fig. 3f) and its ventral surface lacks a conspicuous mat of plumose setae.

The mesial margin of the palm of the chela has a row of 8-9 tubercles (Fig. 4a, b). The lateral margin of the propodus is without spiniform tubercles, and the opposable margin often shows a prominent excision (Fig. 4a). The opposable margin of the propodus carries about 7 solid white denticles and 1 larger and corneous distal one, that is usually spikier and a bit out of line (Fig. 4c). The opposable margin of the dactyl has a row of about 10 solid white denticles, among them a bigger one that is located in the mid-part of the row and out of line (Fig. 4d). The tips of the dactylus and propodus are pointed and corneous (Fig. 4a, c, d). The mesial side of the carpus possesses a group of tubercles and spines (Fig. 4e). The ventral surface of the merus has two rows of more than 10 tubercles and spines each (Fig. 4e), and the basio-ischium has a ventral row of a few tubercles (Fig. 4e). The basis of the cheliped has no mesial spine.

**FIGURE 4.**
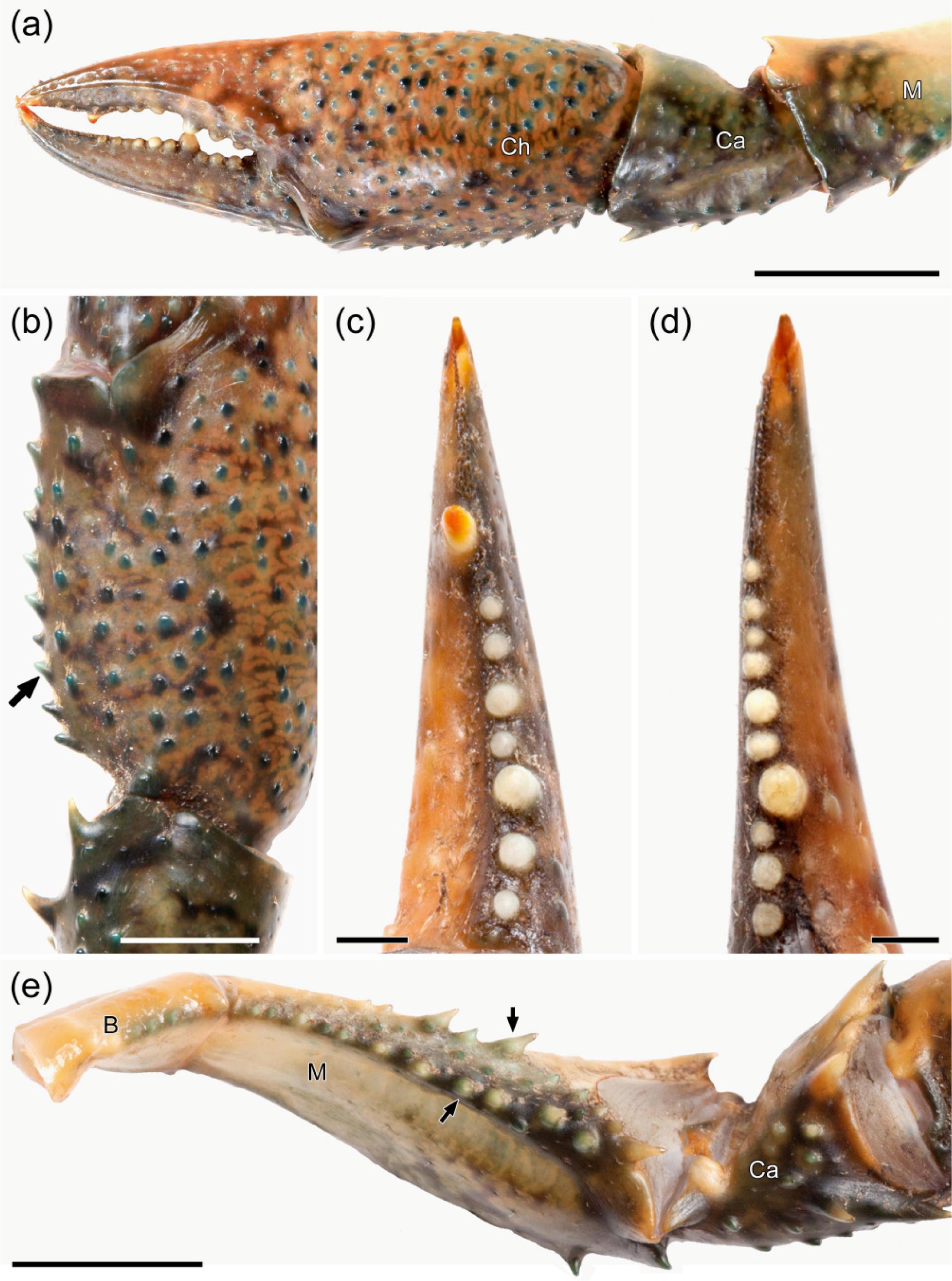
Taxonomically relevant characters on cheliped of marbled crayfish of 9.3 cm TL. Dorsal view of right cheliped. Ch, chela; Ca, carpus; M, merus. Bar: 10 mm. (b) Row of tubercles (arrow) on mesial margin of palm. Bar: 5 mm. (c) Row of denticles on propodus. Bar: 1 mm. (d) Row of denticles on dactylus. Bar: 1 mm. (e) Ventral side of merus with two rows (arrows) of tubercles and spines. B, basio-ischium with row of tubercles. Bar: 5 mm.

The sternum cephalic to the *annulus ventralis* lacks projections or tubercles and is not overhanging the *annulus* (Fig. 5a). The *annulus ventralis* is bellshaped and has an S-shaped sinus (Fig. 5a, d). It is freely movable against the pre-annular sternum and is about 1.6 times as broad as long (Fig. 5a). The post-annular sclerite is dome-shaped and of similar width than the *annulus ventralis* (Fig. 5a).

**FIGURE 5.**
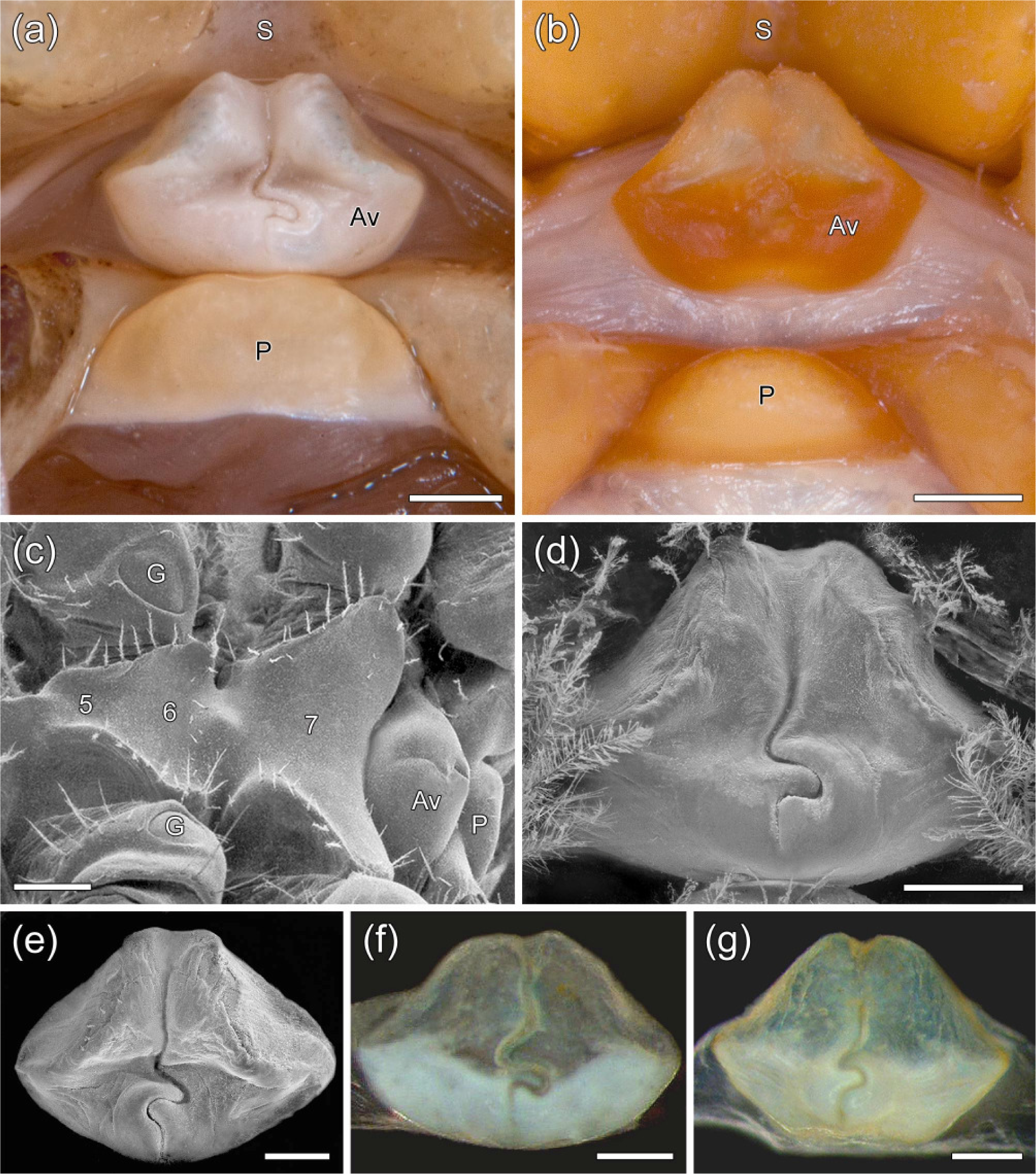
*Annulus ventralis* in marbled crayfish and *Procambarus fallax.* (a) *Annulus ventralis* (Av) and postannular sclerite (P) in fresh marbled crayfish of 9.3 cm TL from Lake Moosweiher. The *annulus* is bell-shaped, has a medial depression and is bisected by a trough leading caudally into an S-shaped sinus. S, pre-annular sternal plate. Bar: 1 mm. (b) *Annulus ventralis* and postannular sclerite in ethanol-fixed *P. fallax* of 6.9 cm TL from the Everglades. Bar: 1 mm. (c) Development of *annulus ventralis* and postannnular sclerite in marbled crayfish juvenile of 1.6 cm TL. The sternal plates of the 5th to 7th thoracomeres (5-7) are fused. G, gonopore. Scanning electron micrograph. Bar: 0.2 mm. (d) Scanning electron micrograph of *annulus ventralis* of laboratory-raised first-time spawning marbled crayfish of cm TL. Bar: 0.5 mm. (e) Scanning electron micrograph of *annulus ventralis* of repeatedly spawned marbled crayfish of 6.8 cm TL. Bar: 0.5 mm. (f) *Annulus ventralis* from exuvia of laboratory-raised marbled crayfish of 5.5 cm TL. Bar: 0.5 mm. (g) *Annulus ventralis* from exuvia of laboratory-raised *P. fallax* of 4.8 cm TL. Bar: 0.5 mm. (c-e) from Vogt et al., 2004, and (f, g) from Vogt et al., 2015.

The pleon is narrower than the carapace (Fig. 2a). The pleura have rounded ventral margins (Fig. 1a, 2b). The first pleopods are present. The telson has mediolateral groups of spines (Fig. 3d). The outer ramus of the uropods has a median horizontal row of spines and the inner ramus has a longitudinal ridge with tubercles terminating in a spine that is not reaching the posterior end of the ramus (Fig. 3d). The diameter of the pleopodal eggs is ~1.5 mm.

The *annulus ventralis* (Fig. 5a-g) or spermatheca is a species-specific trait in the Cambaridae (Hobbs, 1981) and is therefore of particular interest for our comparison between marbled crayfish and *P. fallax.* It is located between the fused sternal plates of the 5th-7th thoracomeres and the postannular sclerite (Fig. 5c). The *annulus* is similar in marbled crayfish (Fig. 5a, d-f) and *P. fallax* (Fig. 5b, g) but deviates in form from the sperm receptacles of other cambarids (Hobbs, 1981, 1989; Vogt et al., 2015). In marbled crayfish and *P. fallax* it is bell shaped, has usually a medial depression and is bisected by a trough leading caudally into an S-shaped sinus (Fig. 5a, b, d-g). In marbled crayfish, the anlage of the *annulus ventralis* is first visible in stage 4 juveniles and is fully developed shortly before first spawning. In adult specimens it is still modified to some degree with increasing body size and age (Fig. 5d, e). There is also certain variability among specimens of the same size. For example, in some individuals there exists a distinct submedian depression, while in others this depression appears to be subdivided by a horizontal ridge. Moreover, the sinus is sometimes reversed. Similar variability of the *annulus ventralis* was also observed in *P. fallax* (this study; Hobbs, 1981).

The most variable morphological characters in marbled crayfish are the rostrum (Fig. 2a, 3a,b, 6a), the areola (Fig. 6a), the denticles on the dactylus and propodus of the chelipeds (Fig. 4c, d), and the spination of the various articles of the chelipeds (Fig. 4a, b, e). The same also holds for *P. fallax* (Hobbs, 1981). In marbled crayfish the length and form of the rostrum can differ considerably among communally raised specimens of the same age. Usually, it is straight (Fig. 3a, b) but can also be slightly flexed vertically and horizontally. In smaller specimens, the anterior tip of the rostrum is mostly spiky but in large specimens it is often blunt. The rostrum usually has two marginal spines in addition to the spiky acumen. These spines and the postorbital, cervical and branchiostegal spines were prominent in all marbled crayfish from Lake Moosweiher (Fig. 3a) and Saxony (Martin et al., 2010b) and all *P. fallax* samples from the wild (this study; Hobbs 1981). In contrast, marbled crayfish from the laboratory populations of G.V. (Fig. 6a) and Gerhard Scholtz (Humboldt University of Berlin; Martin et al., 2010b) had small tubercles instead of spines.

**FIGURE 6.**
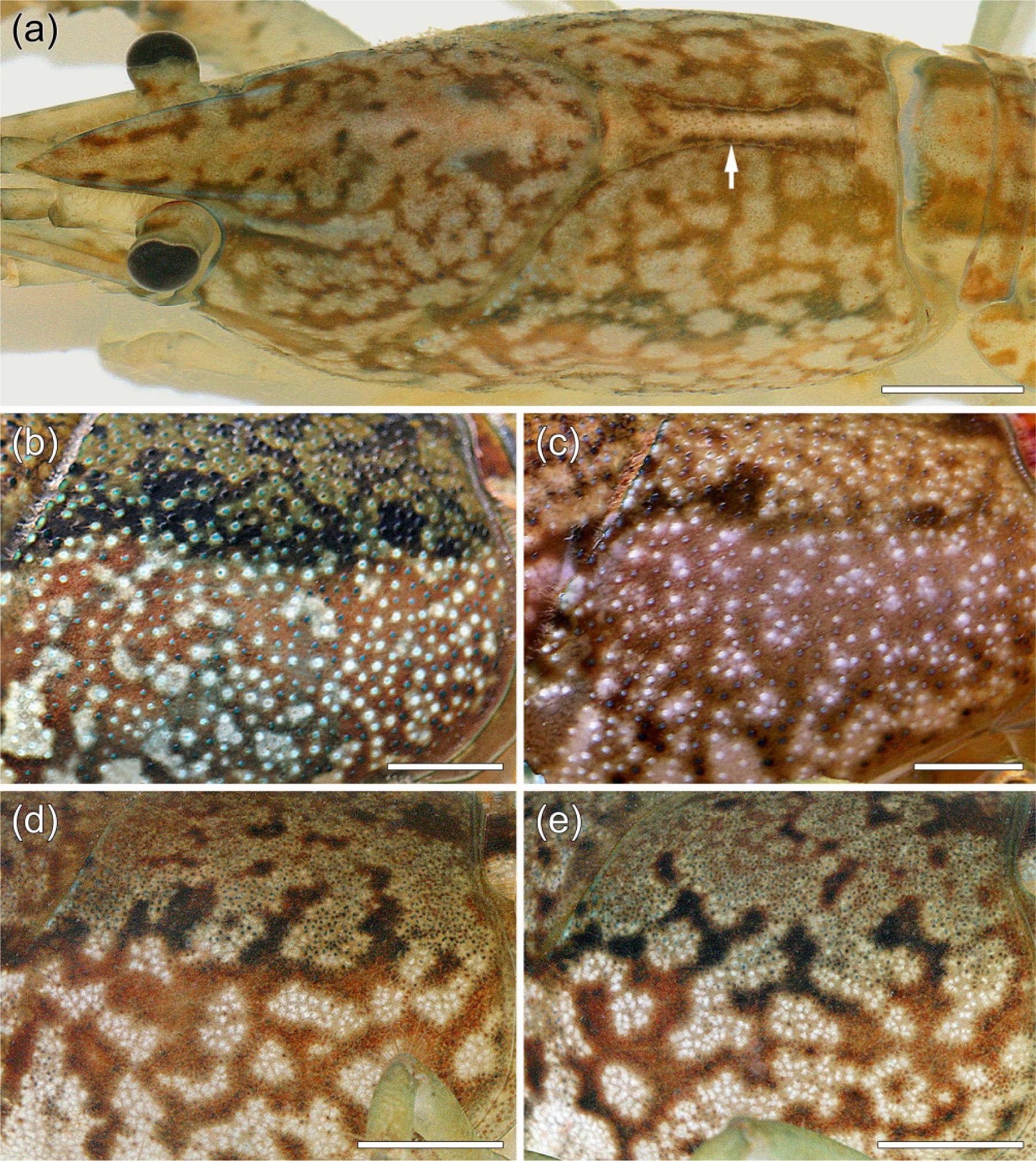
Variation of coloration in marbled crayfish and *Procambarus fallax.* (a) Dorsal view of laboratory-raised marbled crayfish with ocre background coloration. Note differences in marmoration pattern between right and left body sides. Arrow denotes areola. Bar: 5 mm. Banchiostegal part of carapace of marbled crayfish from Lake Moosweiher with relatively dark background coloration. Bar: 5 mm. (c) Branchiostegal carapace of laboratory-raised *P. fallax* with reddish background coloration. Bar: 3 mm. (d, e) Differences of marmoration pattern of branchiostegal carapace of commually reared batch-mates of marbled crayfish. Bars: 3 mm.

The length:width ratio of the areola was 5.59±1.17 (n=20; range: 4.32-7.77) in marbled crayfish and 5.63±1.30 (n=20; range: 4.40-8.81) in *P. fallax.* The denticles on the propodus and dactylus varied in number, size and arrangement. In marbled crayfish, there were 6-9 denticles (n=20) on the propodus (Fig. 4c) and 6-10 denticles (n=20) on the dactylus (Fig. 4d). *P. fallax* had 5-8 denticles (n=20) and 5-9 denticles (n=20), respectively. The number, size and arrangement of the spines and tubercles in the two rows on the ventral side of the merus of the chelipeds (Fig. 4e) were highly variable as well. Spine numbers varied between 10-15 in both rows and both crayfish (n=20 each)

#### 3.1.2 Coloration

Coloration is principally similar in marbled crayfish (Fig. 1a, b, 6b) and *P. fallax* (Fig. 1c, d, 6c) but is highly variable in both crayfish. The background colour of wild specimens of marbled crayfish is often dark brown to olive dorsally (Fig. 2a) and reddish brown laterally (Fig. 2b). There are creamy patches on this darker background, which are most prominent on the branchiostegal part of the carapace (Fig. 1a, b, 2a, b, 6b, d, e). Additionally, there are whitish and black spots, particularly on the pleon and chelipeds (Fig. 2a, b). A median brown stripe is extending dorsally from the rostrum to the caudal margin of the carapace (Fig. 2a). There is a broad black band dorsolaterally on the carapace extending from the cervical groove to the posterior end of the carapace (Fig. 1a, 2b). Each side of the pleon has a prominent undulating black band between terga and pleura (Fig. 1a, 2b) and an additional black band further dorsally, which is either continuous or interrupted. The dorsal sides of the uropods and telson are speckled (Fig. 1a, 3d).

The ventral sides of the cephalothorax and pleon and the pleopods are uniformly creamy. The *annulus ventralis* is more whitish (Fig. 5 a). The pereopods are light brown at their proximal articles becoming increasingly darker distally (Fig. 1a, b, 2b). The dorsal side of the chelipeds displays the same background colour as the body and has lighter blotches and black tubercles (Fig. 1a, 2b, 4a, b). Their ventral side is lighter (Fig. 4e). The ventral sides of the basal articles of the antennule and antenna are also lighter than the dorsal side, but the flagellae are composed of alternating uniformly coloured lighter and darker annuli (Fig. 3a). The scaphocerite is darker dorsally than ventrally and has a dark longitudinal stripe (Fig. 3e).

Variation of coloration in marbled crayfish and *P. fallax* concerns both the background colour and the marbled motifs. In the wild, the background colour is mostly dark brown to olive dorsally and reddish-brown laterally (Fig. 2a, b, 6b), but aquarium specimens have also ochre (Fig. 6a), reddish (Fig. 1b, d, 6c) or bluish background colours. In marbled crayfish, the highest variability of coloration was found for the marbled motifs on the branchiostegal part of the carapace (Fig. 6b, d, e). These motifs are unique in each specimen and become fully established in late adolescents (Vogt et al., 2008). Marmoration as such is genetically determined because there are no uniformly coloured specimens in marbled crayfish and *P. fallax.* In contrast, the marmoration pattern must be epigenetically determined because all marbled crayfish are genetically identical (Martin et al., 2007; Vogt et al., 2008, 2015; Gutekunst et al., 2018). It is the result of stochastic developmental variation (Vogt et al., 2008; Vogt, 2015a), which is best illustrated by pattern differences between right and left body sides of the same individual (Fig. 6a) and comparison of the branchiostegal carapace between communally-raised batch-mates (Fig. 6d, e).

#### 3.1.3 Body proportions

Body proportions were compared between 64 specimens of marbled crayfish of 26.7-101.1 mm TL and 50 females of *P. fallax* of 31.9-88.9 mm TL. Our results revealed that TL:CL and CL:CW were similar between wild populations of marbled crayfish and *P. fallax.* Mean TL:CL of marbled crayfish from Lake Moosweiher was 2.16±0.047 and mean TL:CL of *P. fallax* from the Everglades and the museum collections (originating from different sites in Florida and southern Georgia) was 2.16±0.069 and 2.12±0.048, respectively (Table 4). The coefficient of variance (CV) for TL:CL was 2.2% for marbled crayfish from Lake Moosweiher and 2.3% for *P. fallax* from the NCSM and USNM. The CV for CL:CW was considerably higher, amounting to 5.9% in marbled crayfish from Lake Moosweiher and 10.0% in *P. fallax* from the NCSM and USNM collections, which originated from different geographical regions and habitats.

In marbled crayfish, the body proportions differed markedly between wild and laboratory-raised populations. In this study, we measured a mean TL:CL ratio of 2.16±0.047 for Lake Moosweiher but of 2.26±0.048 for the laboratory colony of G.V. Earlier measurements of 128 marbled crayfish from Lake Moosweiher by Buri (2015) revealed a TL:CL ratio of 2.17, and earlier unpublished measurements of 150 marbled crayfish from the laboratory colony of G.V. revealed a ratio of 2.24, supporting the present data. The laboratory raised specimens were on average more elongate and slender. This may be the result of 15 years of rearing under highly standardized conditions. The 5 marbled crayfish of 4.87-5.14 cm TL from the NCSM collection, which descended from the laboratory colony of Zen Faulkes (University of Texas Rio Grande Valley) had a TL:CL ratio of 2.13±0.026 (r: 2.09-2.16), which is more similar to wild populations.

### 3.2 Similarities and differences in life history traits between marbled crayfish and *Procambarus fallax*

This section includes a meta-analysis of our data and literature data on body size, fertility and longevity of marbled crayfish and *P. fallax.*

#### 3.2.1 Life cycle

The life cycles of marbled crayfish and *P. fallax* occur in fully flooded conditions and consist of embryonic, juvenile and adult periods. The embryos develop in the eggs that are firmly attached to the pleopods by egg stalks. They hatch as juveniles. The first two juvenile stages are lecithotrophic and permanently attached to the pleopods with prominent hooks on their chelae. Stage-3 juveniles and sometimes also stage-4 juveniles adhere most of the time to the maternal pleopods for resting but leave them regularly for feeding. The free juvenile period includes several instars, and the adult life period is characterized by reproductive competence (this study; VanArman, 2003; Vogt & Tolley, 2004; Vogt et al., 2004).

In laboratory-raised marbled crayfish, the duration of embryonic development varies from about 17 to 28 days, depending on water temperature. The postembryonic brooding period lasts 14-25 days depending on living conditions (Vogt & Tolley, 2004; Vogt, 2008a, b). The free juvenile period comprises about 12 juvenile stages and lasts some 4-5 months. The adult period starts with the first reproducing stage, lasts mostly for 1.5-2.5 years and consists of alternating reproduction and growth phases with interspersed moults. Average generation time of marbled crayfish in the laboratory was 6-7 months (Seitz et al., 2005; Vogt, 2008a). The life cycle of *P. fallax* in the laboratory followed principally the scheme of marbled crayfish as observed in the colonies of G.V. and VanArman (2003), but the time spans for the various phases were not yet precisely determined. Embryonic development is about three weeks (VanArman, 2003).

#### 3.2.2 Body size

The mean total lengths of marbled crayfish caught with baited traps and hand fishing nets was 6.7 cm (n=496) in mesotrophic Lake Moosweiher (this study, Chucholl & Pfeiffer, 2010; Günter, 2014; Lehninger, 2014; Wolf, 2014; Buri, 2015), 6.1 cm (n=404) in Lake Šoderica, Croatia (Cvitanić, 2017), 7.0 cm (n=39) in Chorvátske rameno canal, Slovakia (Lipták et al., 2017) and 6.9 cm (n=57) in rice fields and wetlands of Madagascar (Jones et al., 2009). These means correspond to the uppermost range of *P. fallax* sizes caught with dip nets and throw traps in the Everglades (Hendrix et al., 2000; van der Heiden & Dorn, 2017).

We further examined adult sizes of marbled crayfish and *P. fallax* and removed all specimens <18 mm CL (typical juveniles) from the datasets collections of both crayfish. The comparison revealed that specimens larger than 32 mm CL or 7 cm TL are rarely found in *P. fallax* but are frequent in marbled crayfish (Table 5). We have not applied statistics to these data because capture methods and other parameters like trophic status of the habitats were too different to be standardized.

**TABLE 5.** Numbers and percentages of captured crayfish in collections of adult-sized (>18 mm CL) marbled crayfish and *Procambarus fallax* from a variety of freshwater ecosystems.

| **Animals** | **Location** | **>18** | **>28** | **>32** | **>36** | **>40** | **Source** |
| --- | --- | --- | --- | --- | --- | --- | --- |
| Pf<sup>1</sup> | Everglades | 311 | 10 | 1 | 0 | 0 | van der Heiden & Dorn, unpubl. |
| Pf<sup>2</sup> | Everglades | 61 | 9 | 0 | 0 | 0 | Dorn, unpublished |
| Pf | Museums | 39 | 13 | 5 | 4 | 0 | this study |
| Pf | Sum (%) | 100 | 7.8 | 1.5 | 1.0 | 0 |  |
| Mc<sup>3</sup> | Slovakia | 27 | 27 | 27 | 25 | 14 | Lipták et al., 2017 |
| Mc<sup>4</sup> | Madagascar | 58 | 49 | 27 | 11 | 2 | Jones et al., 2009 |
| Mc<sup>3</sup> | Moosweiher | 464 | 385 | 315 | 196 | 81 | this study; Chucholl and Pfeiffer, 2010; Günter, 2014; Lehninger, 2014; Wolf 2014; Buri, 2015 |
| Mc | Sum (%) | 100 | 84.0 | 67.2 | 42.3 | 17.7 |  |

Pf, *Procambarus fallax*; Mc, marbled crayfish. Catching methods: <sup>1</sup> throw trap, <sup>2</sup> baited and unbaited traps, <sup>3</sup> baited trap and hand catch, <sup>4</sup> hand catch. Museum samples were from different habitats in Florida and Georgia, sampling methods unknown.

The maximum size of the 1242 measured marbled crayfish from laboratory colonies and wild populations was 53 mm CL or ~11 cm TL. This largest specimen was from a gravel pit in Reilingen, Germany (n=150; Lyko, 2017). Maximum CLs near 50 mm were determined in oligotrophic-mesotrophic thermal Lake Hévíz, Hungary (50.5 mm CL, n=42; Lőkkös et al., 2016, Lake Moosweiher, Germany (49 mm CL, n=496; this study, Chucholl & Pfeiffer, 2010; Günter, 2014; Lehninger, 2014; Wolf, 2014; Buri, 2015), a Slovakian canal (48.1 mm, n=39; Lipták et al., 2017), and wetlands of Madagascar (47 mm CL, n=57; Jones et al., 2009).

Maximum sizes of *P. fallax* came from unpublished field datasets from the Everglades (n = 35,388; from the laboratories of N.J.D. and Dale Gawlik, Florida Atlantic University, and Joel Trexler, Florida International University), notes on crayfish sizes from wetlands to the northeast of the Everglades and in central Florida (VanArman, 2003; unpublished notes by Peggy VanArman), and large adults recovered from experimental wetlands (unpublished data by N.J.D. from the study Knorp & Dorn, 2014). The maximum size caught from the Everglades sampling was 40.2 mm CL from a naturally nutrient enriched site in the southern Everglades and the second largest was 39.8 mm CL from a nutrient enriched wetland site in the central Everglades (Trexler, unpublished data). The largest *P. fallax* from three wetlands to the north of the Everglades was 39.5 mm CL (VanArman, pers. communication to N.J.D.). From repeated collection in Lake Okeechobee (a eutrophic lake, Glades County, FL) a specimen was recorded at 42 mm CL (David Essian, FAU personal communication to N.J.D.) and specimens from other counties in Florida (e.g., Alachua County, collection of the USNM) measured up to 41 mm CL. Finally, four adults added to the experimental wetlands studied by Knorp & Dorn (2014) were recovered at the end of the summer and all were near 40 mm CL, with the largest being a 41.3 mm CL female.

Scoring all available evidence from various sources, instances of *P. fallax* >32 mm CL or 7 cm TL are quite rare and it seems the maximum size is around 41-42 mm CL, corresponding to ~9 cm TL. In contrast, specimens of marbled crayfish >32 mm CL are frequent and the maximum is around 53 mm CL corresponding to 11 cm TL. The maximum size differences seem incontrovertible as the maximum sized *P. fallax* appear rather uniform and come from multiple counties, multiple wetlands and from some systems with nutrient rich conditions. The larger maximum and typical adult sizes of marbled crayfish may result from faster growth and enhanced longevity. Earlier laboratory experiments suggested that marbled crayfish grow faster than *P. fallax* when kept under identical conditions, but that study was conducted with only a few specimens (Vogt et al., 2015). There is also evidence that marbled crayfish get older and thus can grow longer when compared to *P. fallax* (see 3.2.4), but the database on longevity in *P. fallax* is still too small to make conclusive statements.

#### 3.2.3 Reproduction and fertility

All marbled crayfish from Lake Moosweiher, the laboratory populations of G.V. and N.J.D. the collection of the NCSM and the literature were females (n=1456). These populations exclusively reproduced by apomictic parthenogenesis as shown by the identity of the *COI* sequences and microsatellite patterns in all examined specimens (see 3.3), confirming earlier findings that marbled crayfish is an obligately parthenogenetic all-female crayfish (Martin et al., 2007; Vogt et al., 2008, 2015). In contrast, *P. fallax* reproduces sexually and has males and females in approximately equal ratios. Of 1702 sexed specimens collected by N.J.D. in the Everglades 51.2% were females and 48.8% were males (van der Heiden & Dorn, 2017), and of the 2200 sexed *P. fallax* in the USNM collection 54.8% were females and 45.2% were males (this study). The USNM specimens were collected in different counties and habitats throughout the entire distribution range of *P. fallax* in Florida and southern Georgia.

Pairings of marbled crayfish females and *P. fallax* males performed by G.V. (n=3 pairs) and N.J.D (n=2 pairs) revealed that both crayfish copulated but the offspring was always pure marbled crayfish, as demonstrated by exclusive female sex of the progeny. These data indicate reproductive isolation as already demonstrated by earlier mating experiments (Vogt et al., 2015). Those experiments revealed pure female offspring and identical microsatellite patterns between mothers and offspring. The *P. fallax* males had different microsatellite patterns and did not contribute to the genome of the offspring.

In the laboratory of G.V., marbled crayfish laid eggs in any month of the year except of June but marked peaks of egg-laying were observed around spring and autumn equinoxes (this study; 2015b). First time spawners usually had ages (since hatching) between 150 and 250 days, in an exceptional case 524 days. The latest spawning ever recorded occurred at day 1530 of life (Vogt, 2010). The smallest berried female had a TL of 3.4 cm but usually first time spawners were >4 cm TL. Most females spawned twice a year but some spawned once or three times. The maximum number of clutches per female and lifetime recorded so far was seven (Vogt, 2010). *P. fallax* females in the Everglades had an average size at maturity of 4 cm TL (Hendrix et al., 2000). A long-term experiment with 6 females of *P. fallax* in the laboratory of G.V. revealed one to two spawnings per year and a maximum of three spawnings per lifetime (Vogt, 2018c). The population trends in the Everglades suggest a similar 5-7 month pause between reproductive events (van der Heiden & Dorn, 2017).

In wild marbled crayfish and *P. fallax* populations, berried individuals can be found throughout most of the year. In Lake Moosweiher egg-carrying marbled crayfish were even found in December and January at water temperatures of 5-7°C (Buri, 2015; Christian Günter, personal communication to M.P.). Nevertheless, both crayfish seem to have two major annual recruitment periods in their habitats. In Lake Šoderica, berried marbled crayfish were particularly frequent in June and October/November (Cvitanić, 2017). In Shark Slough, Everglades National Park, Hendrix et al. (2000) found ovigerous *P. fallax* females in every month except July but most ovigerous females were sampled in March and late fall. In the central Everglades, cohorts of small juveniles of *P. fallax* were found in early winter (November-January) with falling water and in early summer (June-July) with rising water (van der Heiden & Dorn, 2017), confirming two major annual recruitment periods per year.

In the marbled crayfish colony of G.V., the lowest number of pleopodal eggs counted at the day of spawning was 51 in a specimen of 15 mm CL. The maximum numbers of pleopodal eggs and juveniles ever reported for marbled crayfish were 731 (Vogt et al., 2015) and 647 (Lipták et al., 2017), respectively. The maximum clutch sizes determined by us from *P. fallax* of the NCSM collection were 135 and 127 eggs in specimens of 25.9 and 27.5 mm CL, respectively. These females were collected from river Styx in Alachua County, FL. The largest clutch of *P. fallax* reported from the Everglades wetlands was 130 eggs from a female of 26 mm CL (Hendrix et al., 2000), and the highest egg counts for a *P. fallax* in the outdoor colonies/experimental systems of N.J.D. were 307 and 333 for females of 28.6 and 33.3 mm CL, respectively.

The fertility (count of pleopodal eggs per female and clutch) of marbled crayfish was statistically compared to fertility of *P. fallax* using all available published and unpublished datasets that the authors could assemble. We digitized egg count and carapace length data from published figures in papers and reports (marbled crayfish: Seitz et al., 2005; Jones et al., 2009; Liptak et al., 2017; *P. fallax:* Hendrix et al., 2000). A few points could not be extracted because they were overlain by other points. Unpublished data from berried marbled crayfish in the laboratory came from G.V. and N.J.D. Most of the *P. fallax* fertility data came from wild-caught animals in Everglades wetlands in the southern third of peninsular Florida. Data collected by N.J.D. and especially from the laboratory of Joel Trexler contributed to the fertility datasets for *P. fallax.* A few individuals were obtained from the collections of the NCSM and USNM. Altogether 215 data points were assembled (124 marbled crayfish and 91 *P. fallax*) (Table 2).

The fertilities of marbled crayfish and *P. fallax* were statistically compared using analysis of covariance. Carapace lengths and egg counts were both transformed with natural logarithms to linearize the relationship between size and fertility and make variances similar between the species. After analyzing the full dataset, four large outliers (i.e., the largest residuals with highest leverage) were removed from the analysis. One outlier was physically impossible, probably representing a data entry error. Removal of the outliers did not change the qualitative conclusions but made differences slightly more conservative. Interaction terms, between egg count and carapace length were non-significant (p > 0.3) and dropped from the final models. The analyses did not meet all linear model assumptions (i.e., residuals were not normally distributed), but inspection of the remaining points with the largest leverage indicated that most of them were making the results conservative. The marbled crayfish dataset included larger females than the *P. fallax* dataset, and so we also ran a second analysis after truncating the datasets to a maximum of 33 mm CL. The results were qualitatively equivalent so that we only report the full analysis.

There was considerable egg count variation for all sizes of marbled crayfish and female *P. fallax.* Statistical analysis of the egg counts indicated that fertility was a function of carapace length (F_1,208_ = 348.8, p < 0.0001) and that after controlling for size marbled crayfish were generally more fertile than *P. fallax* (F_1,208_ = 17.6, p < 0.0001) (Fig. 7). Based on the available data the model-predicted egg counts indicate that, on average, marbled crayfish carry 40-41% more eggs than an equivalent sized female *P. fallax* (Table 6). However, a large fraction of the fertility data for *P. fallax* came from oligotrophic wetlands (i.e., the Everglades), whereas the data for marbled crayfish came from the laboratory or from various water bodies of unknown trophic status (e.g., Jones et al., 2009). Confirmation of the fertility differences will require an experimental examination under controlled nutrient/food availability.

**FIGURE 7.**
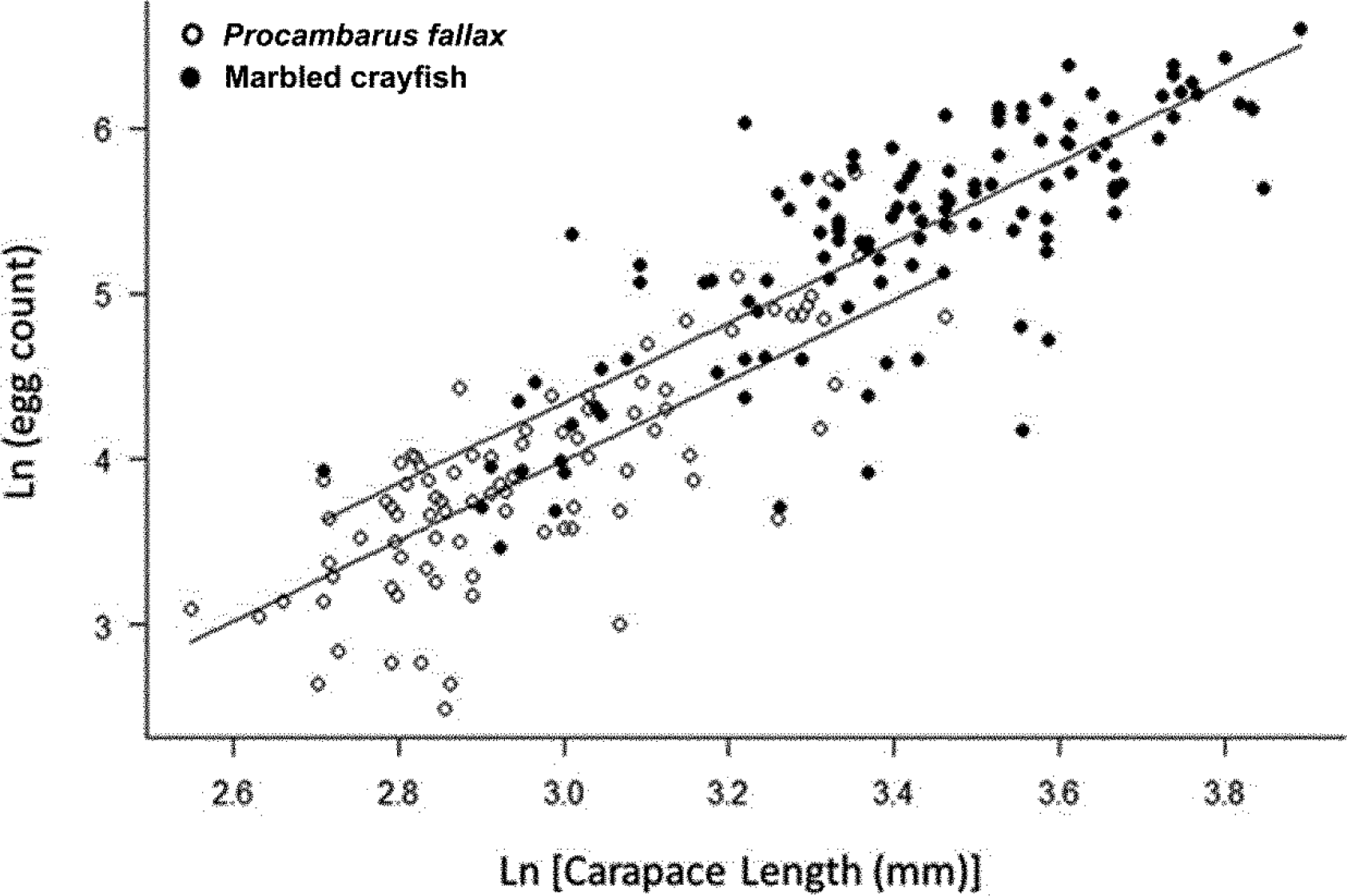
Fertility (eggs per clutch and female) of marbled crayfish and *Procambarus fallax* as a function of female size. The data were pooled from various sources and both variables were plotted on a natural log-transformed scale. The linear model prediction lines were included. The graph shows that fertility is positively correlated with body size in both species but is higher in marbled crayfish of equal sizes.

**TABLE 6.** Model-predicted mean egg counts (± 95% confidence interval) for marbled crayfish and *Procambarus fallax* of various carapace lengths from the linear model analysis.

| Carapace length (mm) | Marbled crayfish | *Procambarus fallax* |
| --- | --- | --- |
| 15 | 38<br>(31-46) | 27<br>(24-30) |
| 20 | 76<br>(67-87) | 54<br>(49-59) |
| 25 | 129<br>(119-143) | 92<br>(82-103) |
| 30 | 202<br>(187-218) | 143<br>(124-165) |

#### 3.2.4 Longevity

Directly measured life span data for marbled crayfish and *P. fallax* are only available for laboratory reared specimens. In the colony of G.V. marbled crayfish that reached the adult stage lived for 718 days on average. Mean life span of the longest lived 10% was 1154 days and maximum longevity was 1610 days (Vogt, 2010). The oldest *P. fallax* raised under the same conditions reached an age of 882 days (this study). However, the number of long-term raised *P. fallax* (n=6) was much lower than of long-term raised marbled crayfish (n=49), which may lead to an underestimation of the maximum life span of *P. fallax.*

Size-frequency distribution data of wild populations suggest that *P. fallax* in wetlands of southern Florida typically live for 1-2 years (Hendrix et al., 2000; van der Heiden & Dorn, 2017), whereas marbled crayfish in Lake Moosweiher typically live for 2-3 years (Wolf, 2014; see 3.4), principally supporting the longevity differences observed in the laboratory. Unfortunately, there are no longevity data available for marbled crayfish in tropical Madagascar and for *P. fallax* in lakes and rivers of northern Florida and southern Georgia, which would make life span comparison between both crayfish more meaningful.

### 3.3 Similarities and differences in genetic features between marbled crayfish and *Procambarus fallax*

In this section we compare the mitochondrial *COI* genes and nuclear microsatellite patterns of our marbled crayfish and *P. fallax* samples and supplement this analysis with literature data on genomic features of both crayfish. Special emphasis is given to intraspecific variability. The *COI* sequences were used for the establishment of a consensus tree on the taxonomic relationship of marbled crayfish with *P. fallax* and other cambarid species.

#### 3.3.1 Mitochondrial *COI* genes and phylogenetic tree

Sequencing of *COI* genes of 4 marbled crayfish from Lake Moosweiher, 3 marbled crayfish from the NCSM and 4 *P. fallax* from the outdoor colony of N.J.D revealed identity of all marbled crayfish and sequence differences to *P. fallax* in 1-4 bases (Genbank accession numbers of our sequences will be provided upon acceptance of the manuscript). Alignment with the Basic Local Alignment Search Tool (BLAST) of *COI* sequences of 26 marbled crayfish from our study and far distant localities in Germany, Italy, Sweden, Madagascar and Japan (sequences from GenBank) confirmed 100% identity (Table 7). In contrast, the *COI* sequences of 13 *P. fallax* from our study, the German aquarium trade and different sites in southern Florida (sequences from GenBank) differed from the marbled crayfish reference sequence from Lake Moosweiher in 1-5 of 658 bases or 0.15-0.76%. Differences in bases were recorded for a total of 10 sequence positions (Table 7). The *COI* sequences from GenBank of *P. leonensis* (Hobbs, 1942) (n=1), *P. seminolae* Hobbs, 1942 (n=1) and *P. lunzi* (Hobbs, 1940) (n=1), the three other species of the Seminolae group, differed from the marbled crayfish reference sequence in 6.11%, 7.44% and 8.76%. Alignment of sequences from GenBank further revealed intra-species variation of 0% in marbled crayfish (n=26), 1.04% in *P. fallax* (n=13), 0.99% in *P. alleni* (Hobbs, 1940) (n=8), 3.19% in *P. paeninsulanus* (Faxon, 1914) (n=20) and 3.83% in *P. clarkii* (Girard, 1852) (n=20). These results confirm genetic identity of all marbled crayfish and demonstrate that the difference between marbled crayfish and *P. fallax* is small enough to fall within the intraspecific range of *P. fallax.*

**TABLE 7.**
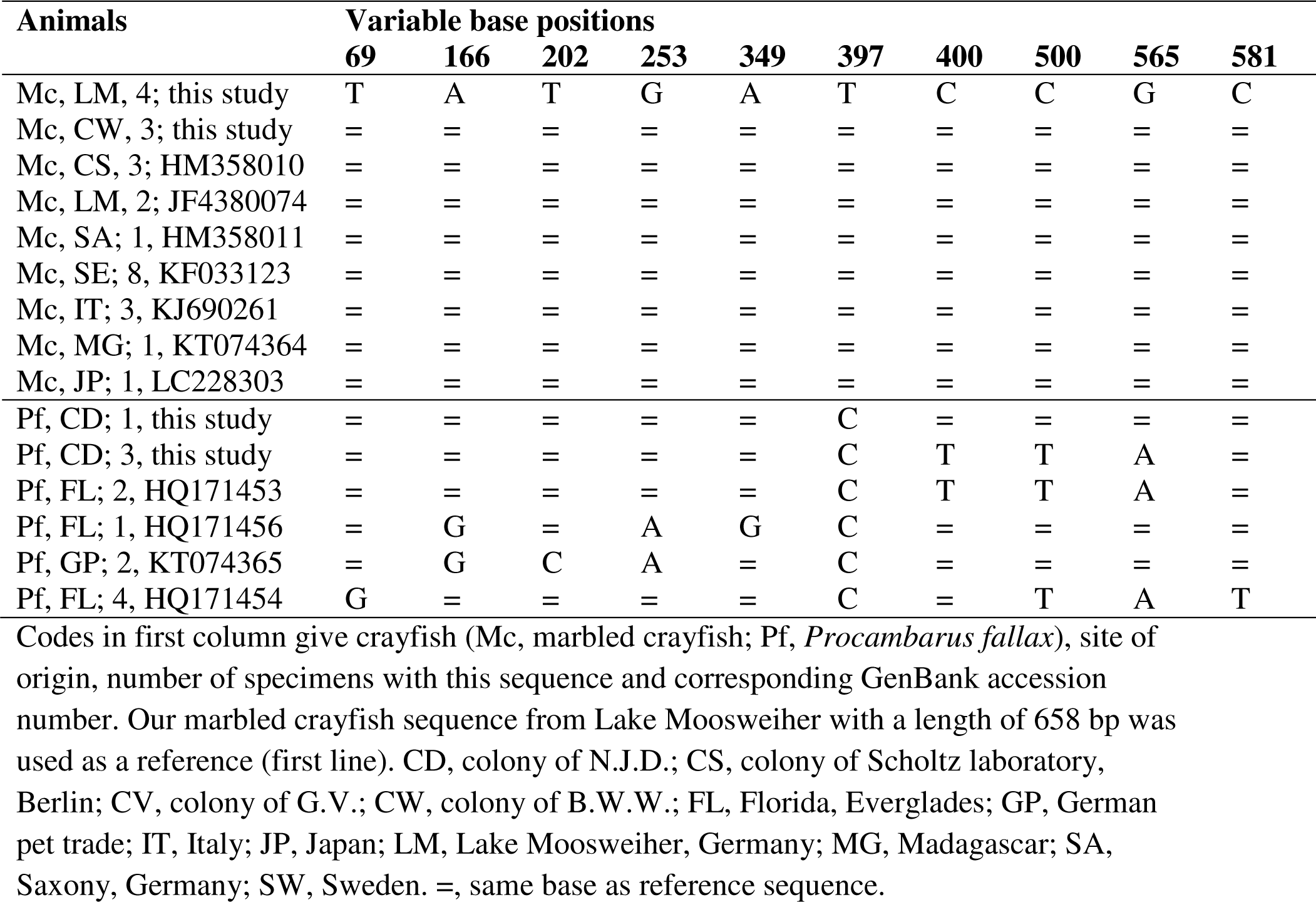
Differences of *COI* sequences between and within marbled crayfish and *Procambarus fallax.*

Earlier analysis of the mitochondrial *12S rRNA* and *COI* genes of marbled crayfish, *P. fallax* and 9 further cambarids revealed closest relationship of marbled crayfish with *P. fallax* (Martin et al., 2010a). The *12S rRNA* genes were even identical between both crayfish. We have now constructed a new consensus tree using the *COI* sequences of marbled crayfish, *P. fallax* and 25 further species of the 440 Cambaridae, among them 15 of the 167 *Procambarus* species (Table 3). Included are for the first time species that were regarded as closest relatives of *P. fallax*, namely *P. seminolae, P. leonensis* and *P. lunzi.* Hobbs (1981) assorted these species together with *P. fallax* in the Seminolae group of the subgenus *Ortmannicus.* We have further included *Procambarus* species that occur sympatrically with *P. fallax* in Florida and southern Georgia (Hobbs, 1942, 1981) and could potentially have hybridized with *P. fallax.* These are *P. seminolae, P. spiculifer* (Le Conte, 1856), *P. paeninsulanus, P. alleni, P. pygmaeus* (Hobbs, 1942) and *P. pubischelae pubischelae* Hobbs, 1942 (Table 3). The tree in Fig. 8 shows a distinct cluster composed of marbled crayfish and *P. fallax*, confirming the close taxonomic relationship of marbled crayfish and *P. fallax.* Marbled crayfish is even nesting within *P. fallax. Procambarus leonensis* and *P. seminolae*, two members of the Seminolae group that was erected on the basis of the male gonopods (Hobbs, 1981) form the sister clade to the *P.* fallax/marbled crayfish cluster. *P. lunzi*, another putative member of the Seminolae group branches off a bit more distantly.

**FIGURE 8.**
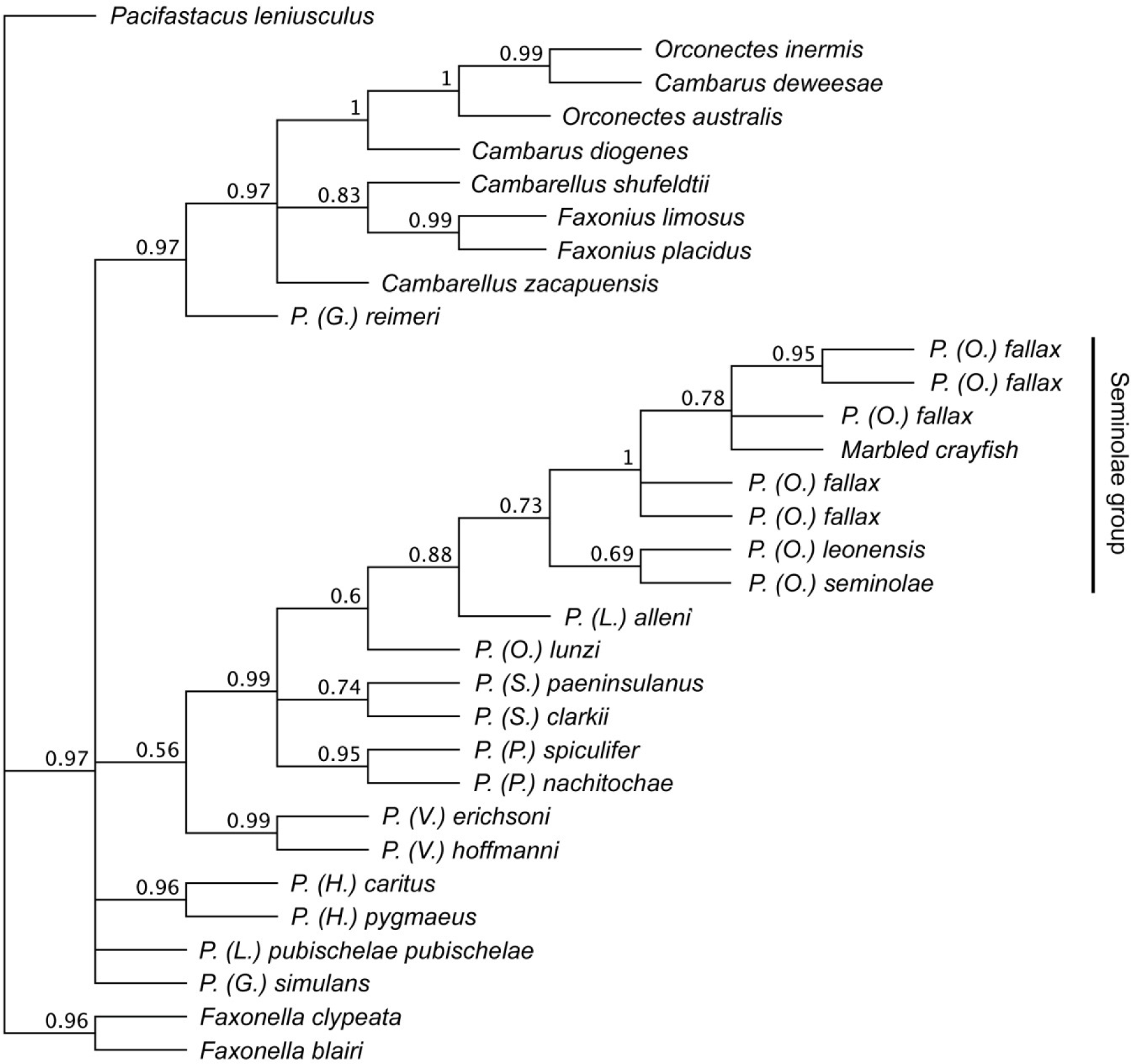
Taxonomic position of marbled crayfish in the Cambaridae. The phylogenetic consensus tree is inferred from 606 bp long fragments of the mitochondrial *cytochrome oxidase subunit I (COI*) genes. Posterior probability values are shown. A clade with a value >0.70 is regarded as well supported. Included are marbled crayfish from Lake Moosweiher, *Procambarus fallax* from various sources, *Procambarus* species from different subgenera (Ortmannicus, Leconticambarus, Scapulicambarus, Pennides, Villalobosus, Hagenides, Girardiella) and species of other genera of the Cambaridae. Marbled crayfish and *P. fallax* form a distinct cluster, and marbled crayfish is even nested within the *P. fallax. Procambarus seminolae and P. leonensis*, two members of the Seminolae group of the genus *Ortmannicus* branch off next to the *P.* fallax/marbled crayfish cluster.

#### 3.3.2 Microsatellites and other nuclear genomic features

The analysis of the microsatellites *PclG-02, PclG-04* and *PclG-26* revealed uniformity in marbled crayfish and heterogeneity in *P. fallax* (Table 8). Our 7 marbled crayfish from Lake Moosweiher and the NCSM had identical allele combinations for *PclG-02* (267 bp/271 bp/303 bp), *PclG-04* (159 bp) and *PclG-26* (189 bp/191 bp). The same pattern was earlier determined for 31 marbled crayfish from German laboratory lineages, Lake Moosweiher and Madagascar (Vogt et al., 2015). In contrast, our 4 *P. fallax* from Florida had di-allelic combinations of the fragments 259 bp, 263 bp, 267 bp and 283 bp for *PclG-02*, mono-allelic 159 bp for *PclG-04* and di-allelic combinations of 169 bp, 185 bp, 191 bp, 193 bp and 197 bp for *PclG-26* (Table 8). The 13 *P. fallax* from a family group investigated earlier (Vogt et al., 2015) had mono-allelic and di-allelic combinations of the fragments 239 bp, 255 bp and 267 bp for *PclG-02*, mono-allelic 159 bp for *PclG-04* and mono-allelic and di-allelic combinations of 179 bp, 185 bp and 207 bp for *PclG-26* (Table 8). Interestingly, 4 of the 6 alleles of marbled crayfish were found among the 17 *P. fallax* examined so far: 267 bp at locus *PclG-2*, 159 bp at *PclG-04*, and 189 bp and 191 bp at *PclG-26.* In contrast, the 4 *P. alleni* and 3 *P. clarkii* investigated by Vogt et al. (2015) had quite different alleles in these three loci (Table 8).

**TABLE 8.** Microsatellite patterns in marbled crayfish, *Procambarus fallax, P. alleni* and *P. clarkii.*

| **Animals** | **PclG-02** | **PclG-04** | **PclG-26** |
| --- | --- | --- | --- |
| Marbled crayfish (n=53) | 267/271/303 | 159 | 189/191 |
| *P. fallax* (n=17) | 239<sup>2</sup> | 159 | 179 |
|  | 239<sup>2</sup> | 159 | 179/185 |
|  | 239/255<sup>2</sup> | 159 | 185 |
|  | 239/255<sup>2</sup> | 159 | 185/207 |
|  | 255<sup>2</sup> | 159 | 185 |
|  | 255<sup>2</sup> | 159 | 179/185 |
|  | 255<sup>2</sup> | 159 | 179/207 |
|  | 255<sup>2</sup> | 159 | 185/207 |
|  | 255/267<sup>2</sup> | 159 | 207 |
|  | 255/267<sup>2</sup> | 159 | 185/207 |
|  | 259/263<sup>1</sup> | 159 | 193/197 |
|  | 259/283<sup>1</sup> | 159 | 185/191 |
|  | 267<sup>2</sup> | 159 | 179/207 |
|  | 267<sup>2</sup> | 159 | 179/207 |
|  | 267<sup>2</sup> | 159 | 185/207 |
|  | 267/283<sup>1</sup> | 159 | 169/189 |
|  | n.d.<sup>1</sup> | 159 | 185/191 |
| *P. alleni* (n=4) | 329/357 | n.d. | n.d. |
|  | 360 | 187 | 183 |
|  | 357/384 | 187 | 161 |
|  | 360 | n.d. | 195 |
| *P. clarkii* (n=3) | n.d. | n.d. | 219 |
|  | 211/218 | n.d. | 219/227 |
|  | 228 | n.d. | 219 |

Given are fragment lengths in base pairs. n.d., not determined. Sources for marbled crayfish: this study; Vogt et al., 2015; Pârvulescu et al., 2017. Sources for *P. fallax*: <sup>1</sup>, this study; <sup>2</sup> Vogt et al., 2015. Sources for *P. alleni* and *P. clarkii*: Vogt et al., 2015.

Investigations by Martin et al. (2016) revealed that marbled crayfish and *P. fallax* have identical haploid sets of 92 chromosomes. However, both crayfish differ markedly with respect to ploidy level, number of chromosomes per body cell, genome size and DNA content per body cell (Table 9). Marbled crayfish are triploid, have 276 chromosomes in non-endopolyploid body cells and have a haploid genome size of 3.5 × 10^9^ base pairs (Vogt et al., 2015; Martin et al., 2016; Gutekunst et al., 2018). In contrast, *P. fallax* are diploid, have 184 chromosomes in normal body cells and possess a haploid genome size of 3.9 × 10^9^ base pairs. Despite of the reduced haploid genome size marbled crayfish include more total DNA per body cell than *P. fallax* because of triploidy.

**TABLE 9.** **Nuclear genomic differences between marbled crayfish and *Procambarus fallax*.**

| Genomic feature | Marbled crayfish | *P. fallax* | Source |
| --- | --- | --- | --- |
| Haploid number of chromosomes | 92 | 92 | Martin et al., 2016 |
| No. of chromosomes per body cell<sup>1</sup> | 276 | 184 | Martin et al., 2016 |
| Haploid genome size (Gbp) | 3.5 | 3.9 | Gutekunst et al., 2018 |
| DNA content per body cell (Gbp)<sup>1</sup> | 10.5 | 7.8 | calculated from ploidy level and genome size |
| Global DNA methylation (%) | 2.4 | 2.9 | Vogt et al., 2015 |

Gbp, 10<sup>9</sup> base pairs; <sup>1</sup> non-endopolyploid cells.

Marbled crayfish have on average ~20% lower global DNA methylation levels than *P. fallax* (Table 9) (Vogt et al., 2015). Methylation of cytosines is an important epigenetic modification of the DNA involved in gene regulation, generation of phenotypic variation and environmental adaptation of animals (Jaenisch & Bird, 2003; Lyko et al., 2010; Vogt, 2017). A more detailed investigation of the genome-wide methylomes by Falckenhayn (2016) revealed differences of cytosine methylation in 240 of 6303 screened genes (3.79%) between a marbled crayfish from the colony of G.V. and a *P. fallax* female from the German pet trade. This difference is almost 10 times higher than the 0.43% difference between two *P. fallax* individuals.

### 3.4 Similarities and differences in behaviours between marbled crayfish and *Procambarus fallax*

In the rearing systems of G.V., marbled crayfish and *P. fallax* laid laterally on the water surface and propelled a mixture of water and atmospheric air through the gill chamber when oxygen in the water was low. Berried females even left the water completely and fanned their clutch in atmospheric air for a few minutes. These behaviours enable both crayfish to survive in waters of very low oxygen concentration as long as they can reach the water surface.

Brooding behaviours were very similar in marbled crayfish and *P. fallax.* The berried females of both crayfish hided in shelters, ceased feeding and were more aggressive than unberried females. They removed the decaying eggs and cleaned and aerated the eggs and juveniles regularly.

Our laboratory-reared marbled crayfish and *P. fallax* showed the typical spectrum of agonistic behaviours of crayfish (defined in Lundberg, 2004). Marbled crayfish have been anecdotally characterized as having low levels of aggression, which could mitigate their potential to compete with other species. However, Jimenez & Faulkes (2011) demonstrated that marbled crayfish can compete with red swamp crayfish *Procambarus clarkii*, which itself is a highly successful invasive species. Such studies are not available for *P. fallax* but since it is one of the most common crayfish species in Florida (Hobbs, 1942; Crandall, 2010) and lives together with often bigger-sized crayfish species it must possess a similarly effective arsenal of agonistic behaviours. Males of *P. fallax* are generally more aggressive than equally sized females and their larger chelae give them an advantage in fights.

In the laboratory, marbled crayfish and *P. fallax* established social hierarchies when reared in groups (this study; Vogt et al., 2008; Farca Luna et al., 2009). The dominants showed aggressive behaviours and the subordinates showed evading behaviours. This is not surprising for *P. fallax*, which have two sexes and are genetically diverse, but for marbled crayfish, which have only one sex and are genetically identical. In marbled crayfish, the biggest female in a group was usually the dominant. In *P. fallax*, the males were dominant over females of equal or moderately larger size. In marbled crayfish, juveniles displayed initially no agonistic behaviours and started to establish social hierarchies from about juvenile stage 7 when their claws became suitable for fighting (Vogt et al., 2008). When a dominant marbled crayfish was removed from his group and grouped together with considerably larger specimens the dominant behaviour was rapidly replaced by subordinate behaviour.

### 3.5 Similarities and differences in ecological features between marbled crayfish and *Procambarus fallax*

The ecological similarities and differences between marbled crayfish and *P. fallax* are summarized in Table 10. Both crayfish are similar with respect to habitats, pH of the water bodies, activity and burrowing but differ with respect to the inhabited climate zones, population structure and invasiveness.

**TABLE 10.**
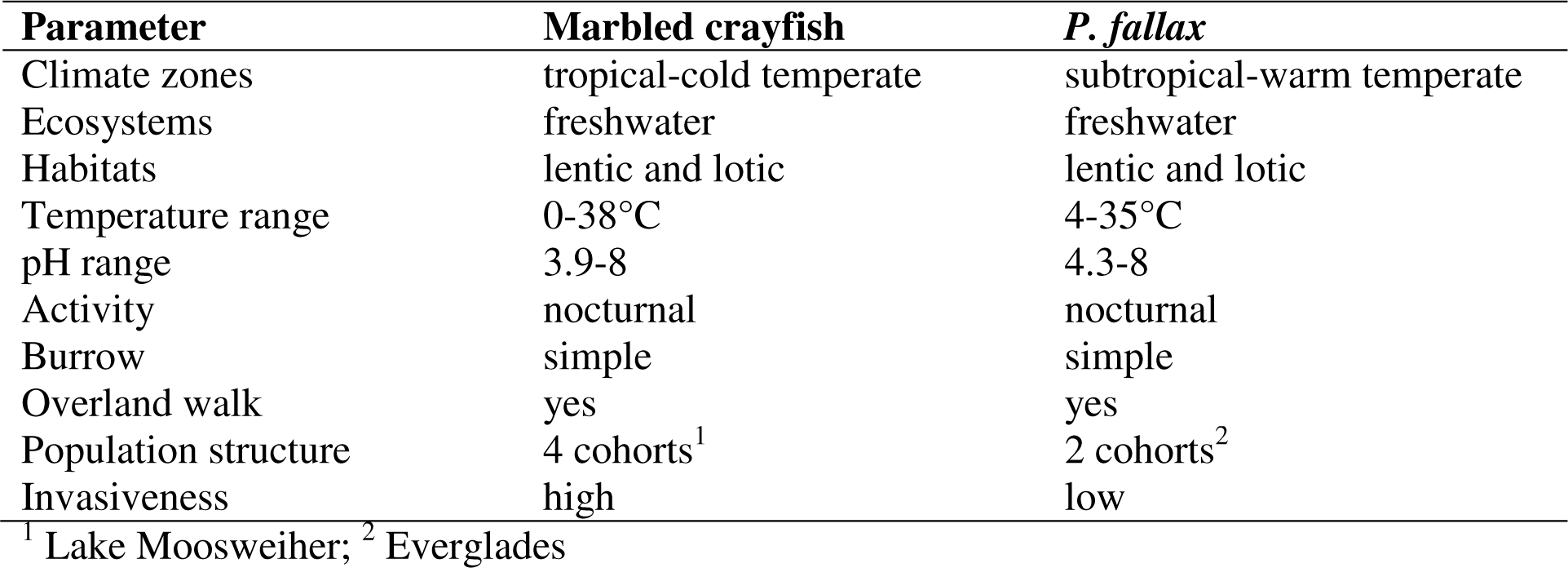
Recorded ecological conditions and population characteristics of marbled crayfish and *Procambarus fallax.*

| Parameter | Marbled crayfish | *P. fallax* |
| --- | --- | --- |
| Climate zones | tropical-cold temperate | subtropical-warm temperate |
| Ecosystems | freshwater | freshwater |
| Habitats | lentic and lotic | lentic and lotic |
| Temperature range | 0-38°C | 4-35°C |
| pH range | 3.9-8 | 4.3-8 |
| Activity | nocturnal | nocturnal |
| Burrow | simple | simple |
| Overland walk | yes | yes |
| Population structure | 4 cohorts<sup>1</sup> | 2 cohorts<sup>2</sup> |
| Invasiveness | high | low |

<sup>1</sup> Lake Moosweiher; <sup>2</sup> Everglades

Throughout Florida and southern Georgia, *P. fallax* is found in lake littoral habitats, ditches, wetlands and slow-moving streams ranging from oligotrophic to eutrophic and with pH ranging from slightly alkaline (pH 8) to acidic (pH 4.2-4.3, measured by N.J.D.) (Hobbs, 1942, 1981; Hendrix et al., 2000; Dorn & Volin, 2009; van der Heiden & Dorn, 2017; Manteuffel-Ross et al., 2018). The presence of vegetation is crucial for their occurrence (Hobbs, 1981; Manteuffel-Ross et al., 2018). Marbled crayfish were found in oligotrophic to eutrophic lakes, brooks and rivers, swamps and rice fields, and acidic (pH 3.9) and polluted waters (Jones et al., 2009; Kawai et al., 2009; Chucholl et al., 2012; Dümpelmann & Bonacker, 2012; Bohman et al., 2013; Gutekunst et al., 2018).

Marbled crayfish and *P. fallax* can walk over land and dig simple burrows (tertiary burrowers). Both crayfish probably do not burrow by preference but when the water table is lowered in the dry season (Hobbs, 1981; Dorn & Volin 2009; Jones et al., 2009; Chucholl et al., 2012; van der Heiden, 2012). Marbled crayfish and *P. fallax* were not yet found in brackish water. However, laboratory experiments revealed that marbled crayfish can survive salt concentration up to 18 ppt for more than 80 days but growth and reproduction stops at 6 ppt (Veselý et al., 2017).

Both crayfish were found in water bodies of variable temperatures. The subtropical wetland habitats of *P. fallax* in Shark Slough of Everglades National Park, southern Florida, had water temperatures of 25°C in spring and fall, peaking at extremes of 35°C in summer and 10°C in winter (Hendrix et al., 2000). In spring-fed Silver River in north central Florida, mean water temperature was 23°C with little variation between seasons (Manteuffel-Ross et al., 2018), and in Lake Okeechobee in south central Florida, temperature fluctuates between 30°C in August and 14°C in January. There are no water temperature data available for the northernmost habitats of *P. fallax* but winter temperatures in Okeefenokee swamp in southern Georgia can fall to about 4°C. The warmest habitat of marbled crayfish reported so far is thermal Lake Hévíz in Hungary with water temperatures of 38°C in summer and 22°C in winter (Lokkös et al., 2016). Water temperature in Antanarivo in the central highlands of Madagascar is about 28°C in summer and 22.5°C in winter. Lake Moosweiher has summer temperatures of 26°C and winter temperatures of 5°C (Chucholl & Pfeiffer, 2010). The coldest habitat so far reported is river Märstaån in Sweden, which has only 0-2°C in winter (Bohman et al., 2013). Marbled crayfish were seen crawling on the ground in this cold water. It is unknown, whether *P. fallax* could survive in such cold waters.

Size-frequency distribution analysis of populations of *P. fallax* in the Everglades revealed two cohorts per year and some bigger specimens suggesting an average longevity of about one year (van der Heiden & Dorn, 2017). Some individuals may live up to two years. In the marbled crayfish population of Lake Moosweiher, Wolf (2014) has found 4 cohorts in late summer/early autumn and some larger specimens. The 4 cohorts probably represent two different years of reproduction, This interpretation is confirmed by peaking of the CLs at 17, 26, 33 and 40 mm, which roughly corresponds with laboratory raised specimens of 6, 12, 18 and 24 months (Vogt, 2010). These data suggest an average longevity of 2 years, and some specimens may live for three years. In the laboratory, most adults reached an age of 2-3 years (Vogt, 2010).

Tracking with transmitters in their natural habitat revealed that the activity of *P. fallax* was considerably higher from 6 pm to 6 am than from 6 am to 6 pm (van der Heiden, 2012). Such tracking studies are not available for marbled crayfish but occasional daily searches in Lake Moosweiher suggest that activity is largely confined to the dark period as well. In the laboratory, both crayfish are mainly active at night. Interestingly, they usually moult during the day in the open arena of the aquarium, when their conspecifics are in the shelters and rest.

*P. fallax* often lives sympatrically with other crayfish species such as *P. alleni, P. seminolae, P. spiculifer, P. paeninsulanus, P. pygmaeus* and *P. pubischelae pubischela* (Hobbs 1942, 1981). Ecological relationships are only known for the pair *P. fallax* and *P. alleni* in the Everglades (Hendrix et al., 2000; Dorn & Trexler 2007; van der Heiden & Dorn, 2017). During the wet season both crayfish were captured together in flooded ditches, and wetlands but during the dry season *P. alleni* (the stronger burrower) can be confined to burrows whereas *P. fallax* often migrates downgradient to deeper flooded areas (Cook et al., 2013. In Lake Moosweiher, marbled crayfish lives sympatrically with the spiny-cheek crayfish *Faxonius limosus*, an invasive crayfish that was introduced to Central Europe in 1890. *F. limosus* is native to eastern North America and is thus naturally adapted to temperate climate. It grows to 12 cm TL and has similar egg numbers per clutch as marbled crayfish but reproduces sexually (Ďuriš et al., 2006; Kozák et al., 2006). Both crayfish seem to occur in equal ratios over the year (Chucholl & Pfeiffer, 2010) but *F. limosus* is predominant in spring and marbled crayfish is predominant in autumn (Günter, 2014; Wolf, 2014; Buri, 2015).

Further ecological differences between marbled crayfish and *P. fallax* concern the distribution to higher latitudes and apparent differences in invasiveness. *P. fallax* is restricted to subtropical biomes and warm temperature regions in Florida and southern Georgia up to latitude N31°, whereas marbled crayfish occurs in tropical biomees and warm and cold temperate regions up to latitude N59.6° (Sweden). In tropical Madagascar, it was found in humid, subhumid and sub-arid climate zones including temperate highland regions above 1000 m altitude (Gutekunst et al., 2018). The difference in distribution between marbled crayfish and *P. fallax* is the result of introductions of marbled crayfish and it is presently unknown whether *P. fallax* could persist in such cold environments as well. Chucholl (2016) calculated an almost double Freshwater Invertebrate Invasiveness Scoring Kit score for marbled crayfish when compared to *P. fallax*, making it a high risk species for Central Europe but also for the USA once released into the wild (Feria & Faulkes, 2011). *P. fallax* has apparently not expanded its range in historical times.

### 3.6 Differences in geographic distribution between marbled crayfish and *Procambarus fallax*

*P. fallax* occurs from tributaries of the Satilla and Suwannee rivers in southern Georgia southward through the Florida peninsula (Fig. 9a) (Hobbs, 1942, 1981; Hendrix et al., 2000; Lukhaup, 2003; VanArman, 2003; van der Heiden & Dorn, 2017). Its distribution range covers some 100.000 km. Marbled crayfish occurs in 11 European countries, Madagascar and Japan (Fig. 9b). In Europe, it was found in one site in the Netherlands (Soes & van Eekelen, 2006), Hungary (Lőkkös et al., 2016), Croatia (Cvitanić, 2017), Romania (Pârvulescu et al., 2017) and Estonia (Estonian Research Council, 2018), two sites in Sweden (Bohman et al., 2013), Italy (Marzano et al., 2009; Vojkovská et al., 2014), the Czech Republic (Patoka et al̤ 2016) and the Ukraine (Novitsky & Son, 2016), five sites in Slovakia (Lipták et al., 2016; Liptak et al., 2017) and 16 sites in Germany (Chucholl et al., 2012; Chucholl, 2016; Lyko, 2017). In Japan, it is known from two localities (Kawai, 2017; Usio et al., 2017) and in Madagascar from many sites (Jones et al., 2009; Gutekunst et al., 2018). All these marbled crayfish populations originated from introductions since the year 2000. In Madagascar, marbled crayfish has spread from the initial introduction site near the capital Antananarivo over 100,000^2^ km as the result of a combination of human and active dispersal. This area roughly corresponds in size to the native range of *P. fallax.*

**FIGURE 9.**
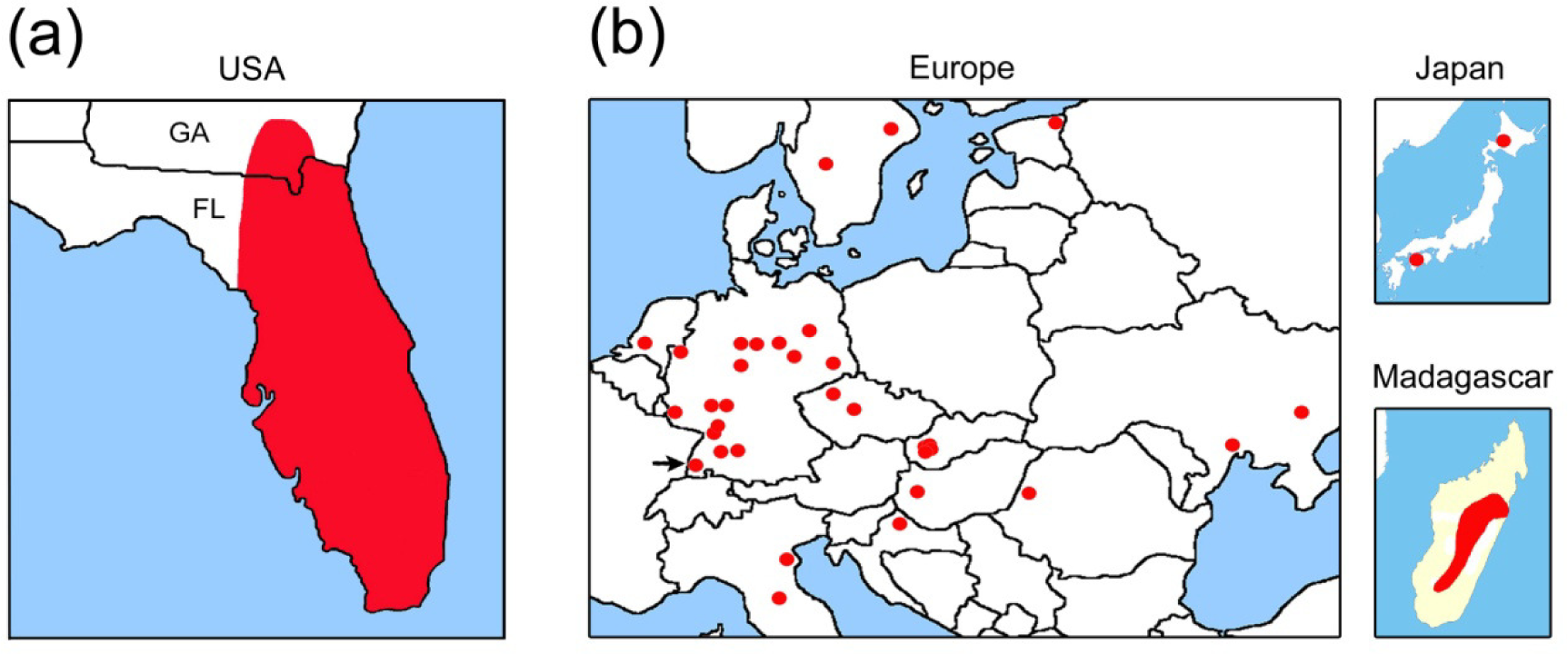
Geographic distribution of marbled crayfish and *Procambarus fallax.* (a) Native range of *P. fallax* in the southeastern USA. The inhabited area is ca. 100.000 km. FL, Florida; GA, Georgia (compiled from data by Hobbs, 1942, 1981; Hendrix et al., 2000; VanArman, 2003; van der Heiden & Dorn, 2017). (b) Populations of marbled crayfish in Europe, Japan and Madagascar. The invaded area in Madagascar is approximately 100.000 km. Arrow denotes Lake Moosweiher (compiled from data by Chucholl, 2016; Lipták et al., 2016, 2017; Lokkös et al., 2016; Novitsky & Son, 2016; Patok et al., 2016; Kawai et al., 2017; Lyko, 2017; Pârvulescu et al., 2017; Usio et al., 2017; Estonian Research Council, 2018; Gutekunst et al., 2018).

There is no evidence in the literature for the occurrence of marbled crayfish in the native range of *P. fallax* (Hobbs, 1942, 1972, 1981; Hendrix et al., 2000; Lukhaup, 2003; VanArman, 2003; van der Heiden, 2012; van der Heiden & Dorn, 2017). Feria & Faulkes (2011) predicted with climate and habitat based Species Distribution Models that marbled crayfish could inhabit a considerably larger geographical area than *P. fallax* when released in the southern states of the USA.

### 3.7 Differences in public policy between marbled crayfish and *Procambarus fallax*

Despite of the unsettled species status, marbled crayfish and *P. fallax* were already treated as different biological and evolutionary units by public policy, which is reflected by different common names and legal regulations.

Common English names of *Procambarus fallax* are slough crayfish and deceitful crayfish. There are no common names in other languages. For marbled crayfish we found the following common names in the literature and internet (in alphabetical order): 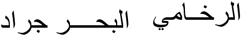 - jarad albahr alrakhamii (Arabian), 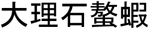 -dàlĭishí áo xia (Chinese), rak mramorový (Czech), mramorni rak (Croatian), marmerkreeft (Dutch), marbled crayfish (English), marmorumita kankro (Esperanto), marmorvähk (Estonian), marmorirapu (Finnish), écrevisse marbrée (French), Marmorkrebs (German), márványrák (Hungarian), gamberi d’acqua dolce marmorizzato (Italian), 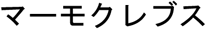 - mamokurebusu (Japanese), 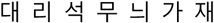 - daeliseogmunuigajae (Korean), foza orana (Malagassy), marmorkreps (Norwegian), rak marmurkowy (Polish), lagosta mármore (Portuguese), racul marmorat (Romanian), MpaMopHbiñ paK - mramornyy rak (Russian), rak mramorovaný (Slovak), marmornati škarjar (Slovenian), cangrejo mármol (Spanish), marmorkräfta (Swedish) and MapMypoBHH paK - marmurovyy rak (Ukrainian). This multitude of common names reflects the worldwide attention to this unusual crayfish.

The legal regulations for marbled crayfish and *P. fallax* are quite different. In the IUCN Red List of Threatened Species *P. fallax* is assessed as “least concern” and is given a NatureServe Global Heritage status rank of G5 (Crandall, 2010). In contrast, as an invasive species marbled crayfish is not subject to conservation regulations. It is rather listed among the invasive alien species of EU concern according to EU Regulation No. 1143/2014 (European Parliament and Council of the European Union, 2014; European Commission, 2017). This regulation prohibits keeping and breeding of marbled crayfish within the Union, placing on the market and releasing into the environment. Keeping for scientific purposes is allowed. Although marbled crayfish has not been found in the wild in the USA there are some legal state regulations due to its availability in the American pet trade (Faulkes, 2015). For example, Missouri has added marbled crayfish to the prohibited species list and Tennessee designated it as “Class V wildlife,” meaning it can only be kept by zoos (Faulkes, 2018).

### 3.8 Discrimination of marbled crayfish and *Procambarus fallax*

The determination of cambarid species is based on the male gonopods. There are no elaborate keys for the determination of females. However, Hobbs (1981) provided a short key to identify females of the genus *Procambarus.* Here, we extend this key to marbled crayfish and *P. fallax.*

1. Presence of annulus ventralis …………………………………………………………… Cambaridae, 2 Absence of annulus ventralis ………………………………………………. Astacidae, Parastacidae
2. Annulus ventralis freely moveable, dactyl of chela distinctly longer than mesial margin of palm, tubercles on mesial surface of latter well developed …………………. *Procambarus*, 3 Otherwise ……………………………………………………………………………………………. other genera
3. Marbled coloration and bell-shaped annulus ventralis ……………. *P. fallax, P. virginalis*, 4 Otherwise ………………………………………………………………………. other *Procambarus* species
4. Males and females present, specimens >7 cm TL rare, maximum size ~9 cm TL, microsatellite *PclG-02* mono-allelic or di-allelic …………………………………………… *P. fallax* Females only, specimens >7 cm TL frequent, maximum size ~11 cm TL, microsatellite *PclG-02* generally tri-allelic ………………………………………………………………… *P. virginalis*

## 4. DISCUSSION

In the following, we examine the taxonomic position of marbled crayfish within the freshwater crayfish family Cambaridae and discuss whether it is justified to consider marbled crayfish as the first asexual species of the almost 15.000 Crustacea Decapoda.

### 4.1 Taxonomic position of marbled crayfish within the Cambaridae

Phylogenetic tree construction with mitochondrial *COI* sequences of marbled crayfish, *P. fallax* and further 25 Cambaridae from different genera clearly revealed that marbled crayfish belongs to the genus *Procambarus* and that it is most closely related to *P. fallax.* The uniform *COI* gene of marbled crayfish differs in less than 1% of bases from the *COI* genes of *P. fallax* but in 6.11%, 7.44% and 8.76% from *P. leonensis, P. seminolae* and *P. lunzi*, the other members of the Seminolae group of the subgenus *Ortmannicus.* The difference in *COI* between marbled crayfish and *P. fallax* is less than the intra-species range of *P. fallax.* This result and the nesting of marbled crayfish within *P. fallax* in *COI* and *12S rRNA* based trees (this study; Martin et al., 2010a) does not argue for raising marbled crayfish to species rank but the data indicate that the maternal ancestor of marbled crayfish was a *P. fallax* female. However, several other parameters analysed by us and discussed below do support considering marbled crayfish as a separate asexual and polyploid species.

### 4.2 Problems associated with the erection of polyploid and asexual species

In the last decade, there was an intense debate on whether polyploids and asexuals like marbled crayfish should be described as new species or kept as cytotypes within their parent species. Soltis et al. (2007) advocated for describing polyploids as separate species when they represent distinct evolutionary units and fulfil the criteria of one or more species concepts, despite morphological similarities with their parent species (for species concepts see Wheeler & Meier, 2000; De Quiroz, 2007; Soltis et al., 2007). They further emphasized that the traditional practice to include morphologically similar cytotypes in one species may be convenient but obscures insights into evolution and speciation processes and hinders conservation.

Plant biologists have meanwhile described numerous polyploid species including autopolyploids (e.g., Eriksson et al., 2017). Barker et al. (2016) estimated that there may be some 50,000 undescribed polyploid species in addition to the 350,000 named species in the flowering plants. Zoologists are more hesitant to place morphologically similar diploids and polyploids in different species but there are already some examples. For instance, Schmid et al. (2015) demonstrated that many of the diploid-polyploid anuran populations are in reality a mixture of diploid and polyploid cryptic species rather than different cytotypes of the same species.

Like polyploids, asexuals do not fit into classical species concepts because these were developed for sexual reproducers (reviewed in Mayr, 1963, 1996; Wheeler & Meier, 2000; Coyne & Orr, 2004). However, Barraclough et al. (2003) emphasized the need to investigate speciation in asexuals to answer central questions of evolution such as the predominance of sexual reproduction in animals. Asexuals are an important component of the biodiversity, particularly in plants (De Meeûs et al., 2007). Well known purely asexual higher animal taxa are the bdelloid rotifers and ostracod crustaceans (Mark Welch et al., 2009; Schön et al., 2009). Further examples of formally described asexual animal species are some mites (Ros et al., 2008) and lizards (MacCulloch et al., 1997). For example, the lizard *Darevskia rostombekowi* is obligately parthenogenetic and monoclonal like marbled crayfish.

### 4.3 Arguments for raising marbled crayfish to species rank

In order to test whether it is justified to consider marbled crayfish as a separate species we have applied to all data the Evolutionary Genetic Species Concept for asexuals (Barraclough et al., 2003; Birky & Barraclough, 2009). This concept requires single origin of the new species, discrete clusters of phenotypes and genotypes in the neo-species and parent species, and reproductive or geographic isolation.

#### 4.3.1 Single origin

Our comparison of the mitochondrial *COI* genes of marbled crayfish from far distant geographical regions such as Germany, Sweden, Italy, Madagascar and Japan revealed that all marbled crayfish have the same sequence. Earlier papers with fewer specimens on *COI* and *12s rRNA* genes (Martin et al., 2010a), *16s rRNA* genes (Jones et al., 2009) and even complete mitochondrial genomes (Vogt et al., 2015) also revealed complete identity of all marbled crayfish examined. The same holds for the nuclear microsatellites *PclG-02, PclG-04* and *PclG-26*, which had identical fragment lengths in marbled crayfish samples from different laboratories and various natural habitats and geographical regions (this study; Reinders, 2012; Vogt et al., 2015; Pârvulescu et al., 2017).

The identity of these commonly used mitochondrial and nuclear markers, which showed considerable variation in *P. fallax* (this study) and other freshwater crayfish species (Hulák et al., 2010; Williams et al., 2010; Da Silva et al., 2011), demonstrate monoclonality of marbled crayfish. Gutekunst et al. (2018) have recently compared the entire genomes of 11 marbled crayfish from Madagascar with the reference genome assembled from a specimen of the colony of G.V. and revealed a total of only 416 single nucleotide variants. The whole-genome difference of individuals to the reference genome was only 128-219 SNVs, which corresponds to the normal mutation rate of animals. These data strongly support single origin of marbled crayfish.

#### 4.3.2 Distinct phenotypic clusters

Marbled crayfish and *P. fallax* are similar with respect to morphological characters but differ markedly with respect to fitness traits. We found no qualitative morphological feature that would unambiguously enable discrimination of the two crayfish. The taxonomically relevant morphological traits (Hobbs, 1972, 1981), coloration and major body proportions vary considerably within both marbled crayfish and *P. fallax*, complicating the recognition of morphological differences. The close morphological similarity does not argue for raising marbled crayfish to species rank but supports the hypothesis that marbled crayfish is an autotriploid of *P. fallax* as earlier suggested (Vogt et al., 2015; Martin et al., 2016). Autopolyploids often look similar to their parent species (Lewis, 1967; Soltis et al., 2007), whereas hybrids often display features of both parent species or intermediate features as shown for the crayfish hybrids *Faxonius rusticus* (Girard, 1852) × *F. propinquus* (Girard, 1852) (Perry et al., 2001) and *F. rusticus* × *F. sanbornii* (Faxon, 1884) (Zuber et al., 2012).

In contrast, other phenotypic features like body size, fertility and longevity differ markedly between marbled crayfish and *P. fallax.* Marbled crayfish reach considerably larger body sizes and produce significantly larger clutch sizes than *P. fallax.* The larger clutch sizes are due to bigger growth (egg number is positively correlated with body size) and ~40% increased fertility in equally sized specimens. A further important factor that contributes to higher fertility of marbled crayfish populations is parthenogenesis. In parthenogenetic marbled crayfish all adults produce eggs whereas in sexually reproducing *P. fallax* only 50% of the adults produce eggs.

Marbled crayfish and *P. fallax* females are similar with respect to brooding behaviour, agonistic behaviour and breathing of atmospheric air at low dissolved oxygen in the water. At least the former two behaviours are rather uniform in the Cambaridae (Gherardi, 2002). Behavioural differences between both crayfish are attributed to the presence of males in *P. fallax* and their absence in marbled crayfish. In *P. fallax* the males are often the dominants in groups whereas in marbled crayfish the largest females are usually the dominants.

Marbled crayfish and *P. fallax* are ecologically similar in as far as they are generalists that inhabit very diverse habitats and can stand harsh and seasonally fluctuating environmental conditions. Both crayfish also show effective survival adaptations to heavy predation pressure and competition with other crayfish species. They differ in population dynamics and invasive potential, which is probably related to differences in life history traits and longevity.

These data indicate that the distinct phenotypic cluster criterion is fulfilled for several traits including body size, fertility, longevity, population structure and invasiveness.

#### 4.3.3 Distinct genetic clusters

There are close similarities in the mitochondrial genomes between marbled crayfish and *P. fallax* but striking differences with respect to some nuclear genomic features, suggesting that the distinct genetic clusters requirement is fulfilled as well. The nuclear genomic differences concern ploidy level, haploid genome size, microsatellite pattern and DNA methylation level. Ploidy is 2N in *P. fallax* and 3N in marbled crayfish, and the haploid genome size is about 10% smaller in marbled crayfish than in *P. fallax* (Vogt et al., 2015; Gutekunst et al., 2018). Loss of DNA during or after polyploidization is not uncommon. For example, in synthetic autopolyploids of the plant *Phlox drummondii* an immediate loss of 17% of total DNA has been recorded followed by a further reduction of up to 25% upon the third generation (Parisod et al., 2010). The degree of DNA loss in marbled crayfish is not trivial because it corresponds to several times the entire genome of the well known genetics model *Drosophila melanogaster.* Even if only non-coding sequences were lost it might have significant consequences for gene regulation via alteration of the chromatin structure.

Marbled crayfish and *P. fallax* also differ markedly with respect to genome-wide DNA methylation, an important epigenetic modification of the DNA, influencing gene regulation, phenotype variation and environmental adaptation (Jaenisch & Bird, 2003; Lyko et al., 2010; Vogt, 2017). The reduction of DNA methylation by 20% in marbled crayfish may have led to pronounced changes of gene expression when compared to *P. fallax.* Together with differences in ploidy and related gene dosage alteration these epigenetic differences may be causative of the enhanced fitness traits and higher invasiveness of marbled crayfish.

#### 4.3.4 Reproductive or geographic isolation

Our present and earlier (Vogt et al., 2015) crossbreeding experiments with marbled crayfish females and *P. fallax* males revealed that both mated readily but the offspring was always pure marbled crayfish, suggesting reproductive isolation. Reproductive isolation is probably due to cytogenetic incompatibility rather than behavioural and mechanical barriers because mating behaviours and the morphology of the sperm receptacles are very similar in marbled crayfish and *P. fallax.*

Marbled crayfish is one of the very rare cases of parthenogenetically reproducing animal triploids. Autotriploidization of the whole genome is not too rare in animals but usually remains undetected and has no consequences at the population level because triploids are mostly sterile. For example, 91 of 5142 progeny of Atlantic salmon *Salmo salar* from Norwegian farms were spontaneous but sterile autotriploids (Glover et al., 2015). Saura et al. (1993) have developed a model that explains how triploidy and parthenogenesis can arise through a single event, resulting in a reproductively isolated new species. The genetic changes that have led to parthenogenesis in marbled crayfish can now be identified by comparison of the already available whole genomes of marbled crayfish and *P. fallax* as outlined in Vogt (2018d).

Analysis of the biogeographic literature on crayfishes in Florida and Georgia (Hagen, 1870; Hobbs, 1942, 1981) revealed no evidence for marbled crayfish in the native range of *P. fallax.* In these references, there are no *P. fallax* females larger than 9 cm mentioned that in reality could be undetected marbled crayfish. Conversely, *P. fallax* is absent from the countries where marbled crayfish has established wild populations as the result of introductions, suggesting geographic isolation.

According to current knowledge, marbled crayfish and *P. fallax* are reproductively and geographically isolated. Since we cannot exclude that marbled crayfish is present somewhere in the extensive wilderness of Florida and southern Georgia the geographic isolation argument remains yet vague. Genetic investigation of historical samples of *P. fallax* from museum collections and new surveys in Florida and Georgia would help to clarify this issue.

#### 4.3.5 Diagnosability

Diagnosability is an important criterion for the erection of a new species. This criterion is fulfilled for marbled crayfish, since it can unambiguously be identified and distinguished from *P. fallax* by the easily measurable tri-allelic microsatellite *PclG-02.* In practice, *COI* was mostly used instead of *PclG-02* to genetically identify marbled crayfish (e.g., Bohman et al., 2013; Lőkkös et al., 2016; Usio et al., 2017). However, strictly speaking, *COI* is not as specific as *PclG-02.* If marbled crayfish has originated from *P. fallax* by autotriploidy in evolutionarily recent times as earlier assumed (Vogt et al., 2015) then diploid female descendants of the maternal ancestor lineage of marbled crayfish may still exist. These specimens would have the same *COI* sequence as marbled crayfish but mono-allelic or di-allelic *PclG-02.*

### 4.4 Arguments against raising marbled crayfish to species rank

Arguments that speak against raising marbled crayfish from "forma" to species rank may come from the close similarity of morphological traits and mitochondrial genes in marbled crayfish and *P. fallax.* However, the morphological similarity is no disqualifier because there are numerous morphologically very similar cryptic animal species described that were only identified by genetic differences or marked life history differences (e.g., Hebert et al., 2004; Bickford et al., 2006; Mills et al., 2017). The similarity of the mitochondrial genomes of marbled crayfish and *P. fallax* is also no disqualifier. It rather supports the hypothesis that marbled crayfish has arisen from a female *P. fallax* in relatively recent evolutionary times.

Colleagues advocating for treating marbled crayfish as a parthenogenetic lineage of *P. fallax* rather than a separate species usually referred to the examples of water flea *Daphnia pulex* and New Zealand mud snail *Potamopyrgus antipodarum*, which have sexual and parthenogenetic populations in a single species. However, the obligately parthenogenetic populations of these species do neither meet the single origin criterion nor the reproductive or geographic isolation criterion of the Evolutionary Genetics Species Concept. In *P. antipodarum* triploid lineages originated many times from sexually reproducing populations and coexist with their sexual ancestors (Neiman et al., 2011). The same holds for *D. pulex*, which includes cyclic parthenogenetic and obligately parthenogenetic populations. The obligate parthenogens have evolved multiple times from the cyclic parthenogens, occur together with them and cross-breed occasionally with them (Hebert & Finston, 2001; Dufresne, 2011).

Vogt et al. (2015) have earlier hypothesized that marbled crayfish may have originated from a single *P. fallax* in the German pet trade in about 1995, because there is no hint in the biogeographic literature on the existence of marbled crayfish in North and Central America, the home of the Cambaridae (Hobbs, 1942, 1972, 1981, 1989). It is well known that the descendants of the first parthenogenetic marbled crayfish in Germany spread via aquarists and the pet trade, first in Germany and later throughout the world. Specimens of this homogeneous world-wide "indoor population" were repeatedly released into the wild, explaining easily the monoclonality of all marbled crayfish populations in Europe, Africa and Asia and their geographic separation from *P. fallax.*

However, the marbled crayfish could also have originated somewhere in the native range of *P. fallax* by spontaneous autotriploidy. An individual of this mutant population could unintentionally have been brought to Germany, where parthenogenesis was detected. Repeated searches for crayfish in the southeastern USA by C.L., who is well familiar with both crayfish, revealed *P. fallax* and many other species but no marbled crayfish (Lukhaup, 2003; unpublished data). However, if, contrary to expectation, evidence should show up that marbled crayfish is present in the native range of *P. fallax* and was there already before 1995, then the single origin and reproductive isolation arguments had to be re-examined by genetic analysis.

### 4.5 Résumé: Marbled crayfish is a distinct evolutionary unit and should be considered as a separate asexual species

If one had only the morphology of small and medium sized specimens of marbled crayfish and *P. fallax* and their mitochondrial genes for decision-making, there would be no reason for treating marbled crayfish as a separate species. However, if one considers monoclonality of marbled crayfish, reproductive isolation in laboratory experiments, considerable differences in fitness traits, nuclear genomic features and some ecological characteristics, and the present knowledge on geographic distribution, then it is justified to assign marbled crayfish and slough crayfish to different species named *Procambarus virginalis* and *Procambarus fallax* as earlier proposed (Vogt et al., 2015; Lyko, 2017). All criteria of the Evolutionary Genetic Species Concept for asexuals and the diagnosability requirement for the erection of a new species are well met. In Madagascar, marbled crayfish has already invaded an area of ~100.000 km^2^ that is comparable in size to the native range of *P. fallax* in the southeastern USA. Thus, marbled crayfish represents a geographically separate and numerically large evolutionary unit that in future will evolve independently from *P. fallax*, supporting the proposed species status. Following this argumentation, marbled crayfish would now be the first asexual and the first polyploid species of the almost 15.000 Crustacea Decapoda.

## ACKNOWLEDGEMENTS

We thank Christian Günter (Freiburg) for providing unpublished information on the life history and ecology of marbled crayfish in Lake Moosweiher and Britta Wahl-Ermel (University of Koblenz-Landau) for sequencing of the marbled crayfish and *P. fallax* samples. We further acknowledge the provisioning of size and fertility data for *P. fallax* by Dale Gawlik (Florida Atlantic University, Boca Raton, FL), Joel Trexler (Florida International University, Miami, FL) and Peggy VanArman (Palm Beach Atlantic University, North Palm Beach, FL). A considerable portion of the *P. fallax* fertility data were collected as part of work funded by Task Agreement #p15AC01258 of Cooperative Agreement #H5000-06-0104 between the Everglades National Park and J. Trexler.

